# Serotonergic neuron-glioma interactions drive high-grade glioma pathophysiology

**DOI:** 10.64898/2025.12.10.693579

**Authors:** Richard Drexler, Belgin Yalçın, Rebecca Mancusi, Abigail Rogers, Kiarash Shamardani, Pamelyn J. Woo, Alexandre Ravel, Samuel Wu, Yahaya A. Yabo, Lisa Steger, Carlos Alberto Oliveira de Biagi-Junior, Costanza Lo Cascio, Robert Malenka, Boris D. Heifets, Mariella G. Filbin, Dieter Henrik Heiland, Karl Deisseroth, Michelle Monje

**Affiliations:** Department of Neurology and Neurological Sciences, Stanford University, Stanford, CA, 94305, USA.; Department of Neurosurgery, University Hospital Erlangen, Friedrich-Alexander University Erlangen Nuremberg, Erlangen, Germany.; Dana-Farber/Boston Children’s Cancer and Blood Disorders Center; Harvard Medical School, Boston, MA, USA.; Nancy Pritzker Laboratory, Department of Psychiatry and Behavioral Sciences, Stanford University, Stanford, CA, USA.; Department of Anesthesiology, Perioperative, and Pain Medicine, Stanford University School of Medicine, Palo Alto, CA, USA.; Department of Neurological Surgery, Northwestern University Feinberg School of Medicine, Chicago, IL, USA.; Department of Psychiatry and Behavioral Sciences, Stanford University, Stanford, CA 94305, USA.; Department of Bioengineering, Stanford University, Stanford, CA 94305, USA.; Howard Hughes Medical Institute, Stanford, CA 94305, USA.

## Abstract

High-grade gliomas are lethal brain cancers that are powerfully regulated by glutamatergic neurons through activity-dependent paracrine factors and functional neuron-to-glioma synapses. Here, we report that serotonergic neurons promote the proliferation of high-grade gliomas throughout the brain. Serotonergic neuronal activity drives circuit-specific increases in high-grade glioma proliferation, calcium transients, and reduced survival. This growth-promoting effect is chiefly mediated by activation of the serotonin (5-hydroxytryptamine; 5HT) receptor 5HT_2A_ on glioma cells. Knock out or pharmacological blockade of 5HT_2A_ receptors in glioma abrogated the glioma growth-promoting effects of serotonergic neuronal activity, while serotonergic psychedelic drugs robustly promote malignant cell proliferation. Gliomas alter serotonergic neuronal activity patterns, resulting in elevated serotonin release into the tumor microenvironment. Together, these findings uncover pathogenic, feed-forward interactions between serotonergic neurons and glioma cells.

## Main Text

High-grade gliomas, including glioblastoma and diffuse midline glioma (DMG), are highly aggressive and typically fatal malignancies, representing the leading cause of death from primary central nervous system tumors(*1*). This grim prognosis is driven by the complex molecular pathogenesis of these tumors, compounded by crucial interactions within the tumor microenvironment. A key feature of these interactions involves the central nervous system itself, particularly neurons(*2*). Neuron-cancer interactions have emerged as central to brain cancer pathophysiology, particularly in gliomas(*3–11*) and brain metastases(*12–14*). Neuronal activity promotes brain cancer pathophysiology through the release of paracrine growth factors(*3*, *4*, *10*, *11*) and through neurotransmitter signaling, including true neuron-to-cancer synapses that depolarize glioma cell membranes, activating intracellular calcium signaling and promoting malignant cell proliferation and migration(*5–9*). Previous studies have demonstrated that various neuronal inputs—including glutamatergic(*5*, *6*, *8*), GABAergic(*7*), and cholinergic(*15*, *16*) neurons—robustly influence glioma growth and progression. Not only do neurons promote brain cancer growth, but brain cancer cells remodel functional neural circuits(*17*) and cause neuronal hyperexcitability (*5*, *18*, *19*) to further augment neuronal inputs to the tumor.

While the roles of some neuronal subpopulations in glioma progression have been explored, the contributions of serotonergic brainstem neurons—primarily located in the raphe nuclei, which are subdivided into the dorsal raphe nucleus (DR) and median raphe nucleus (MR)—remain understudied. Given the widely distributed projections of serotonergic neurons in the brain - influencing a broad range of physiological and behavioral functions in both health and disease - and the predilection for glioma formation within and spread to the brainstem where serotonergic neuronal soma reside, we hypothesized a potential role for 5HT neurons in glioma pathophysiology.

Here, we investigate the influence of serotonergic neurons and serotonin (5-hydroxytryptamine; 5HT) on glioma pathophysiology in a range of neuroanatomical locations and report that dorsal and median raphe serotonergic long-range projections promote a robust, circuit-specific increase in glioma growth. The effects of serotonergic neurons on glioma cells are mediated by both known (e.g. NLGN3) and previously underappreciated growth factors such as ICAM-1 and NCAM-1, and through activation of the serotonin receptor HTR2A. At the same time, gliomas cause a progressive dysregulation of serotonergic neurons that increases serotonin release into the tumor microenvironment and is associated with behavioral changes such as decreased active coping.

### Serotonergic Neurons Promote Glioma Growth

In order to test whether serotonergic neurons and the consequent release of serotonin (5HT) promotes glioma pathophysiology, we developed optogenetic models to control serotonergic neuronal activity in mice bearing genetically accurate murine or patient-derived human gliomas. First, we utilized an immunocompetent SERT-Cre mouse model crossbred with a TIGRE2.0 transgenic mouse expressing the floxed opsin ChRmine to achieve expression of the opsin ChRmine in serotonergic neurons specifically (SERT-Cre_+/wt_ x Ai230_flx/wt_; ***fig. S1A-C***). Genetically engineered murine H3K27M-mutant DMG cells were allografted to the dorsal raphe nucleus (DR) or median raphe nucleus (MR) – the main sources of 5HT neuronal cell bodies – and optogenetic stimulation (20 Hz, ten 5-ms pulses of light delivery every 3 seconds over 30 minutes) was performed after four weeks of tumor engraftment (***Fig. 1A***). Glioma cells infiltrated extensively into both nuclei (***Fig. 1B***), and stimulation of either nucleus drove the proliferation of glioma cells located in or near the raphe nuclei (***Fig. 1C***).

**Fig. 1:**
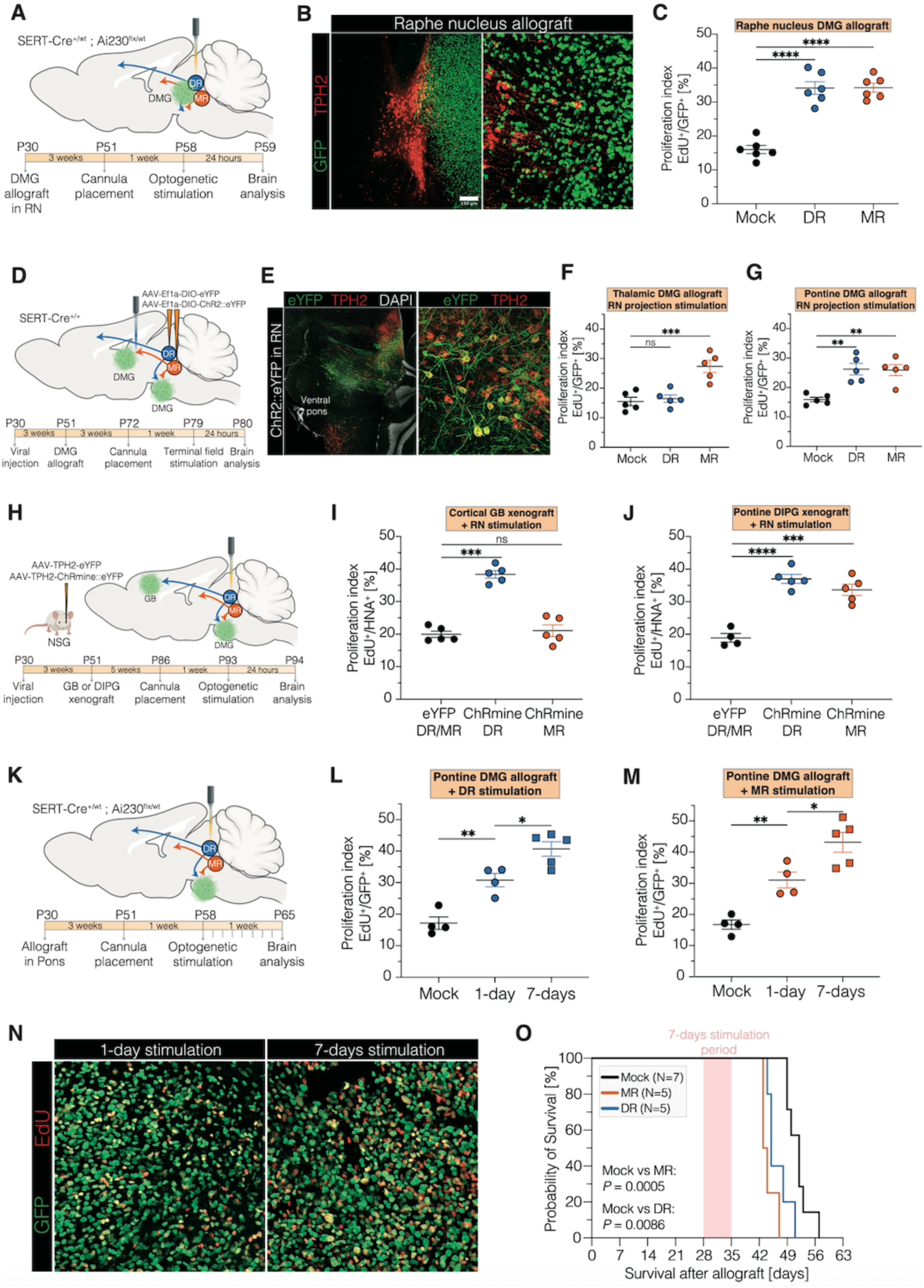
Circuit-specific effects of DR^5HT^ and MR^5HT^ neurons on glioma. **A.** Schematic of experimental paradigm for optogenetic stimulation of DR^5HT^ and MR^5HT^ neurons in mice bearing H3K27M DMG. Four-week-old SERT-Cre_+/wt_ x Ai230_flx/wt_ mice (P28-30) were allografted with a H3K27M MADR tumor model adjacent to the raphe nucleus, with optic ferrule placement into the DR or MR three weeks after allografting. Optogenetic stimulation for 30 minutes of the DR or MR was performed four weeks after allografting, followed by perfusion 24 hours after stimulation. **B.** Confocal micrographs showing GFP_+_ tumor cells adjacent to the raphe nucleus. GFP: green, TPH2: red, Aq: aqueductus. **C.** Proliferation index (EdU_+_/GFP_+_) of raphe nucleus allografts in mice either stimulated in DR or MR or mock-stimulated (“Mock”) (Mock, DR, and MR, n=6 mice/group). Unpaired two-tailed Welch’s t-test; ****p < 0.0001. Data=mean ± SEM. **D.** Schematic of the experimental paradigm for optogenetic stimulation of DR^5HT^ and MR^5HT^ projections in mice bearing H3K27M DMG in the ventral pons and thalamus. Four-week-old SERT-Cre_+/+_ mice (P28–30) were virally injected with AAV-DJ-EF1α-DIO-eYFP or AAV-DJ-EF1α-DIO-hChR2(H134R)::eYFP in the DR or MR, followed by allograft of H3K27M MADR tumor cells into the ventral pons or thalamus three weeks later. Optic ferrule placement into the thalamic or ventral pontine glioma microenvironment was performed three weeks after allografting. Optogenetic stimulation was conducted for 30 minutes, four weeks after allografting, followed by perfusion 24 hours after stimulation. **E.** Representative confocal micrographs showing viral labeling (AAV-DJ-EF1α-DIO-hChR2(H134R)::eYFP) of MR^5HT^ neurons in a sagittal section. eYFP: green, TPH2: red, DAPI: white. **F.** Proliferation index (EdU_+_/GFP_+_) of thalamic allografts in mice either stimulated of DR^5HT^ or MR^5HT^ projections or mock-stimulated (“Mock”) (Mock, DR, and MR, n=5 mice/group). Unpaired two-tailed Welch’s t-test; ***p < 0.001, ns: non-significant. Data=mean ± SEM. **G.** Proliferation index (EdU_+_/GFP_+_) of ventral pontine allografts in mice either stimulated of DR^5HT^ or MR^5HT^ projections or mock-stimulated (“Mock”) (Mock, DR, and MR, n=5 mice/group). Unpaired two-tailed Welch’s t-test; **p < 0.01. Data=mean ± SEM. **H.** Schematic of experimental paradigm for xenografting with optogenetic stimulation of either DR or MR in immunodeficient mice. Four-week-old NSG mice (P28-30) were retroorbitally injected with rAAVPHP.eB-Tph2::eYFP or rAAVPHP.eB-Tph2::ChRmine-eYFP and xenografted into the ventral pons with patient-derived DMG cells (SU-DIPG17) or into the M2 cortex with patient-derived glioblastoma cells (SF232) three weeks after viral vector delivery. After five weeks of glioma growth, optical ferrules were placed either in the DR or MR and optogenetic stimulation was performed one week later. Mice were perfused 24 hours after optogenetic stimulation. **I.** Proliferation index (EdU_+_/HNA_+_) of M2 cortical xenografts in mice either stimulated in DR (“ChRmine DR”) or MR (“ChRmine MR”) or mock-stimulated (“eYFP”) (Mock, DR, and MR, n=5 mice/group). Unpaired two-tailed Welch’s t-test; ***p < 0.001, ns: non-significant. Data=mean ± SEM. **J.** Proliferation index (EdU_+_/HNA_+_) of ventral pontine xenografts in mice either stimulated in DR (“ChRmine DR”) or MR (“ChRmine MR”) or mock-stimulated (“eYFP”) (Mock, DR, and MR, n=5 mice/group). Unpaired two-tailed Welch’s t-test; ***p < 0.001, ****p < 0.0001. Data=mean ± SEM. **K.** Schematic of experimental paradigm for optogenetic stimulation of DR^5HT^ and MR^5HT^ neurons in mice bearing H3K27M DMG. Four-week-old SERT-Cre_+/wt_ x Ai230_flx/wt_ mice (P28-30) were allografted with a H3K27M MADR tumor model into ventral pons, with optic ferrule placement into the DR or MR three weeks after allografting. Optogenetic stimulation for either 30 minutes for one-day or 10 minutes for 7 straight days of the DR or MR was performed four weeks after allografting. **L.** Proliferation index (EdU⁺/GFP⁺) of ventral pontine allografts in mice stimulated in the DR for 1 day (“1-day”), 7 days (“7-days”), or mock-stimulated (“Mock”) (Mock, 1-day, n=4 mice/group; 7-days, n=5 mice). Unpaired two-tailed Welch’s t-test; **p < 0.01, *p < 0.05. Data=mean ± SEM. **M.** Proliferation index (EdU⁺/GFP⁺) of ventral pontine allografts in mice stimulated in the MR for 1 day (“1-day”), 7 days (“7-days”), or mock-stimulated (“Mock”) (Mock, 1-day, n=4 mice/group; 7-days, n=5 mice). Unpaired two-tailed Welch’s t-test; **p < 0.01, *p < 0.05. Data=mean ± SEM. **N.** Confocal micrographs showing proliferating GFP_+_ DMG cells in ventral pons in 1-day-stimulated (left image) and 7-days-stimulated (right image) mice. GFP: green, EdU: red. **O.** Survival curves of ventral pontine allografts (MADR) in mice treated with either mock stimulation, 1-day stimulation, or 7-day stimulation. Log-rank test.

Having established that serotonergic neuronal activity promotes glioma growth in the raphe nuclei, we next explored the effects of serotonergic neuronal long-range projections on glioma in the thalamus and ventral pons, sites where DMG most commonly originates. The dorsal raphe nucleus projects to the pons and the cortex, while the median raphe nucleus projects to the pons and thalamus(*20*) (***Fig. 1D***). SERT-Cre_+/+_ mice were injected with an AAV carrying floxed channelrhodopsin (ChR2) into the dorsal and median raphe to achieve expression of ChR2 in serotonergic axonal projections (***Fig. 1D-E***, ***fig. S1E***), and DMG cells were allografted into the thalamus or ventral pons. Viral labeling of dorsal and median raphe serotonergic projections revealed prominent presence of serotonergic axons within the ventral pontine glioma microenvironment (***fig. S1D***). Terminal field stimulation of either dorsal or median raphe serotonergic axonal projections in the thalamic or ventral pontine glioma microenvironment resulted in robustly increased glioma proliferation in response to serotonergic axonal activity in a circuit-specific manner (***Fig. 1F-G***), similar to previous observations for the effects of cholinergic midbrain neurons on glioma(*21*). Long-range projections from both nuclei promoted ventral pontine glioma cell proliferation (**Fig. 1G**), while thalamic glioma cells only responded to thalamus-projecting median raphe serotonergic projections (***Fig. 1F***). The response of glioma cells to dorsal or median raphe serotonergic neuronal activity was further validated using patient-derived adult IDH-wildtype glioblastoma and pediatric H3K27M DMG xenograft models in immunodeficient NSG mice, which revealed cortical glioblastoma proliferation only with cortex-projecting dorsal raphe serotonergic neuronal stimulation (***Fig. 1H-I***, ***fig. S1F-G***), and ventral pontine DMG proliferation in response to serotonergic neuronal stimulation of either dorsal or median raphe, as both project to the ventral pons (***Fig. 1J***, ***fig. S1H***). This serotonergic nuclei-dependent, circuit-specific effect on glioma cell proliferation was further validated using the aforementioned transgenic mouse model for optogenetic control of serotonergic neuronal activity (SERT-Cre_+/wt_ x Ai230_flx/wt_). Murine DMG cells were allografted to the ventral pons, thalamus, or neocortex and optogenetic stimulation of dorsal or median raphe serotonergic neurons was performed (***fig. S2A-B***). Consistent with expected axonal projections of dorsal and median raphe, serotonergic neuronal activity from the median raphe drove glioma cell proliferation in the ventral pons and thalamus, but not in cortex, while ventral pontine and cortical – but not thalamic – glioma allografts responded to dorsal raphe activity (***fig. S2C-E***). The circuit-dependent effects reflect the expected projection patterns of dorsal and median raphe serotonergic neurons, which were confirmed by viral tracing of serotonergic axons from median and dorsal raphe in the cortex, thalamus, and ventral pons (***fig. S2F***). Taken together, these findings position serotonergic neuronal activity as another important neuronal driver of glioma growth, alongside glutamatergic(*3*, *5*, *6*, *8*, *10*), GABAergic(*7*), and cholinergic(*15*, *16*, *21*) neuronal activity.

To assess the effects of serotonergic neuronal activity on glioma growth over time, we stimulated dorsal or median raphe serotonergic neurons daily (20 Hz, ten 5-ms pulses of light delivery every 3 seconds over 10 minutes) for 7 consecutive days (***Fig. 1K***). Comparing effects of serotonergic neuronal stimulation once vs daily for 7 days by administering the thymidine analogue EdU at the time of optogenetic stimulation on day 1 (1-day paradigm) or on day 7 (7-day paradigm), we found that daily stimulation for 7 days resulted in a higher glioma proliferation rate compared to a 1 day of stimulation (***Fig. 1L-N***), suggesting a degree of cumulative effects. In mice bearing ventral pontine DMG allografts, daily stimulation over 7 days during the middle of the disease course reduced overall survival (**Fig. 1O**).

The formation of neuron-to-glioma networks facilitates neuronal activity-induced depolarization and consequent calcium transients of glioma cells, a crucial driver of proliferation and growth(*5*, *6*, *8*). To further understand if serotonergic neuronal activity affects glioma calcium transients, we performed fiber photometry of patient-derived glioma cells expressing a genetically encoded calcium indicator (GCaMP6s) - with DMG xenografted to the thalamus or glioblastoma xenografted to the cortex – at the same time as optogenetic stimulation of dorsal or median raphe serotonergic neurons in the brainstem (***Fig. 2A-B***). We found that serotonergic neuronal activity resulted in an elevation in glioma calcium transients in the expected circuit-specific manner, with dorsal raphe serotonergic neuronal stimulation increasing calcium transients in cortical glioma xenografts (***Fig. 2C***), and median raphe serotonergic neuronal stimulation increasing calcium transients in thalamic glioma xenografts (***Fig. 2D***).

**Fig. 2:**
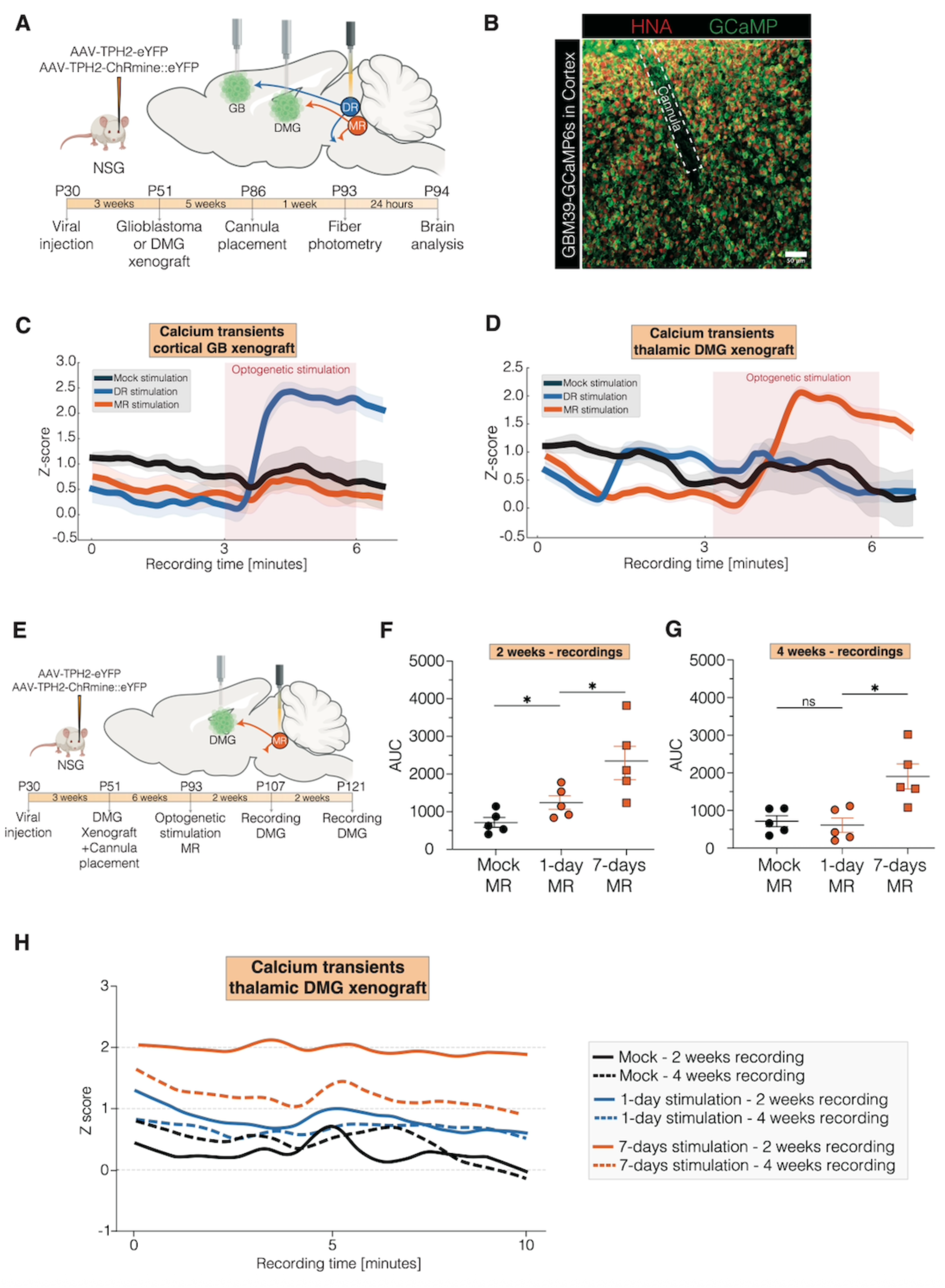
DR^5HT^ and MR^5HT^ neuronal activity triggers long-lasting calcium transients in glioma. **A.** Schematic of the experimental paradigm for fiber photometry recordings of calcium activity in cortical and thalamic xenografts. Four-week-old NSG mice (P28-30) were retroorbitally injected with rAAVPHP.eB-Tph2::eYFP or rAAVPHP.eB-Tph2::ChRmine-eYFP and xenografted into the M2 cortex with patient-derived glioblastoma cells expressing GCaMP6s (GBM39-GCaMP6s) or thalamus with patient-derived DMG cells expressing GCaMP6s (SU-DIPG6-GCaMP6s) three weeks after viral vector delivery. Optical ferrules were placed in the DR or MR, as well as into the xenograft. Fiber photometry recording with simultaneous optogenetic stimulation was performed six weeks after tumor growth. **B.** Confocal micrographs showing M2 cortical xenografts (left image) and fiber photometry cannula placement within the cortical xenograft (right image). GCaMP: green, HNA: red, DAPI: white, scale bars = 1000µm (left image) and 50µm (right image). **C.** Fiber photometry recordings showing averaged GCaMP-labeled calcium transients in cortical glioblastoma cells with simultaneous mock (n=4 mice), DR (n=6 mice), and MR (n=6 mice) stimulation. **D.** Fiber photometry recordings showing averaged GCaMP-labeled calcium transients in thalamic DMG cells with simultaneous mock (n=5 mice), DR (n=4 mice), and MR (n=6 mice) stimulation. **E.** Schematic of the experimental paradigm for fiber photometry recordings of calcium activity in thalamic xenografts. Four-week-old NSG mice (P28-30) were retroorbitally injected with rAAVPHP.eB-Tph2::eYFP or rAAVPHP.eB-Tph2::ChRmine-eYFP and xenografted into the thalamus with patient-derived DMG cells expressing GCaMP6s (SU-DIPG6-GCaMP6s) three weeks after viral vector delivery. Optical ferrules were placed in the MR, as well as into the thalamic xenograft. Mice were stimulated for either 1 day or 7 days, and fiber photometry recordings were performed two and four weeks after the last stimulation session. **F.** Area under the curve of fiber photometry recordings from thalamic xenografts two weeks after stimulation (Mock, 7-days, n=5 mice/group; 1-day, n=4 mice). Unpaired two-tailed Welch’s t-test; *p < 0.05. Data are presented as mean ± SEM. **G.** Area under the curve of fiber photometry recordings from thalamic xenografts four weeks after stimulation (Mock, 7-days, n=5 mice/group; 1-day, n=4 mice). Unpaired two-tailed Welch’s t-test; *p < 0.05, ns: non-significant. Data are presented as mean ± SEM. **H.** Averaged GCaMP fiber photometry traces of thalamic xenografts in mice either mock-stimulated, 1-day-stimulated or 7-days-stimulated (Mock and 7-days, n=5 mice; 1-day, n=4 mice).

To investigate the prolonged effects of 5HT neuronal activity on glioma calcium transients, we employed fiber photometry to measure calcium transients in GCaMP6s-expressing glioma cells within thalamic xenografts (***Fig. 2E***). Calcium transients in thalamic glioma were measured at 2 weeks and 4 weeks after 1-day or 7-days stimulation of median raphe serotonergic neurons. While one day (one session) of stimulation increased calcium transients in thalamic glioma cells compared to mock controls (identically manipulated but without light) during the first recording session, this activity normalized after 4 weeks (***Fig. 2F-H***). In contrast, 7 days of stimulation resulted in persistently elevated calcium transients even after 4 weeks (***Fig. 2F-H***), highlighting the lasting impact of chronic exposure of glioma cells to serotonergic neuronal activity.

### Serotonergic Neuronal Activity–Dependent Paracrine Factors in Brainstem

Given the robust and sustained effects of dorsal raphe and median raphe serotonergic neuronal activity on both glioma cells and neuronal activity in the glioma microenvironment, we next sought to identify the secreted factor profile released as a result of serotonergic neuronal activity. To assess for activity-regulated secreted factors in brainstem, we prepared acute hindbrain slices containing the raphe nuclei from glioma-bearing SERT-Cre_+/wt_ x Ai230_flx/wt_ mice (***Fig. 3A-B***), enabling *ex vivo* optogenetic stimulation, or mock-stimulation (identical manipulation but without light), of ChRmine-oScarlet-expressing serotonergic neurons (***fig. S3A***), followed by collection of conditioned media (CM). Successful serotonergic neuronal stimulation was confirmed by the evident cFos increase in serotonergic raphe nuclei neurons following optogenetic stimulation in this acute slice paradigm (***fig. S3B-C***). Comparing the effects of CM from stimulated slices with mock-stimulated slices enables identification of serotonergic neuronal activity-dependent secreted factors enriched in stimulated CM. As previously demonstrated with CM from forebrain slices (*3*) and retinal+optic nerve explants (*11*), CM collected from stimulated hindbrain slices increased glioma proliferation (***Fig. 3C***, ***fig. S3D***) and migration (***fig. S3E-F***) compared to CM from mock-stimulated controls.

**Fig. 3:**
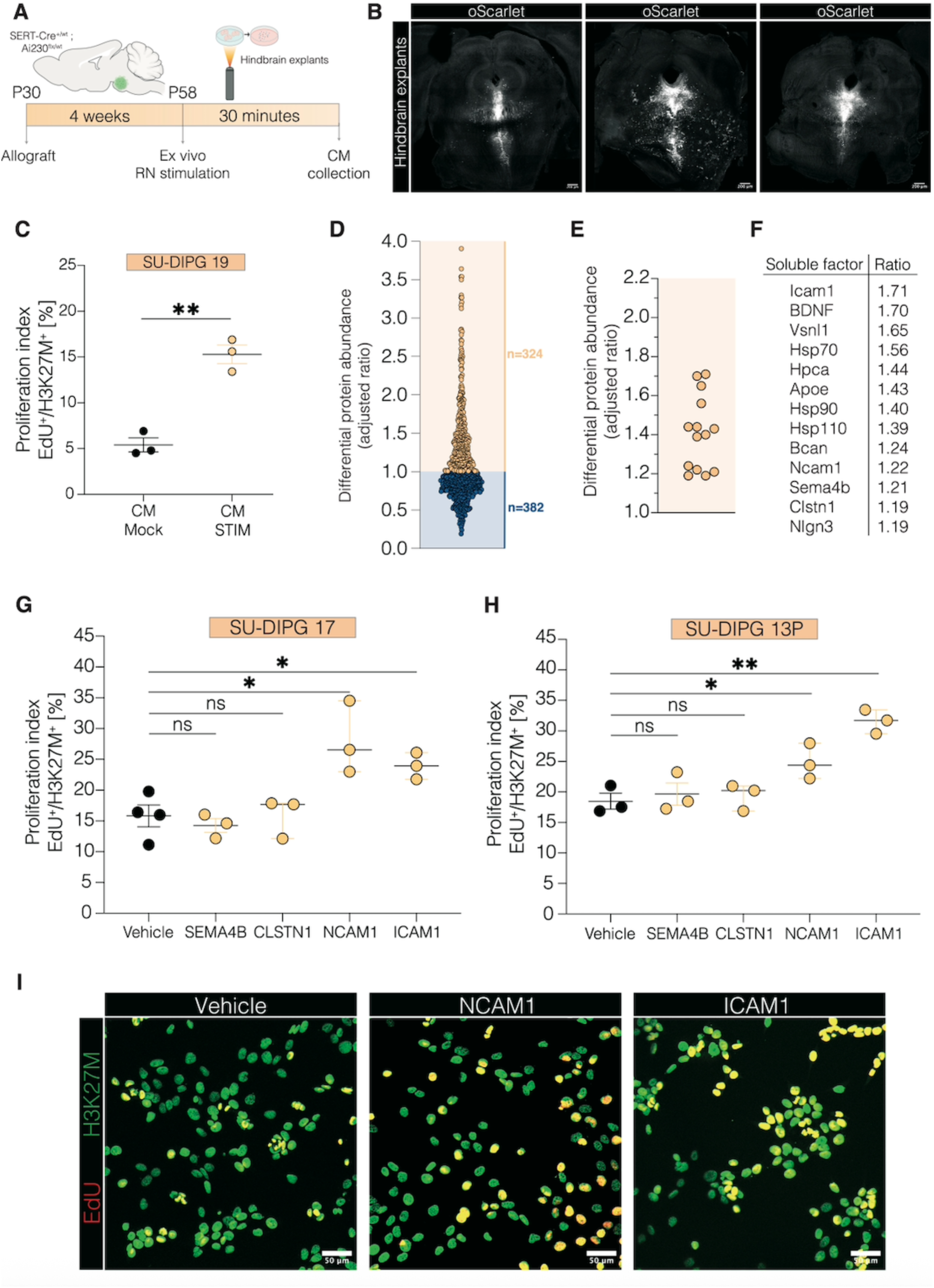
Serotonergic neuronal activity–dependent paracrine growth factors. **A.** Schematic of experimental paradigm for collection of conditioned media (CM) after *ex vivo* optogenetic stimulation of 5HT neuronal cell bodies within the RN in hindbrain explants of 4-week-old SERT-Cre_+/wt_ x Ai230_flx/wt_ mice. **B.** Confocal micrographs of fixed slices used for *ex vivo* optogenetic stimulation. oScarlet: white, scale bars = 200µm. **C.** Quantification of DMG cell proliferation (EdU_+_/DAPI_+_) when adding CM after *ex vivo* stimulation of RN compared to CM from mock-stimulated slices. Unpaired two-tailed Welch’s t-test; **p < 0.01. Data=mean ± SEM; n=three independent experiments, each with three wells per condition; each data point represents the mean of three wells per condition for a given experiment. **D.** Plot illustrating up- and downregulated proteins in conditioned media (CM) following *ex vivo* RN stimulation compared to mock-stimulated slices from glioma-bearing mice, based on LC/MS analysis. **E.** Proteins identified as upregulated after RN stimulation with potential to act as soluble factors. **F.** List of upregulated proteins identified as candidate mediators at the 5HT neuron–glioma interface. **G.** Quantification of DMG cell proliferation (SU-DIPG17) upon addition of recombinant proteins (identified in panel F) to the culture media. One-way analysis of variance (ANOVA) with Tukey’s post hoc analysis; *p* < 0.05, ns: not significant. Data are presented as mean ± SEM; *n* = 3 independent experiments, each with three wells per condition. Each data point represents the mean of three wells per experiment. **H.** Quantification of DMG cell proliferation (SU-DIPG13P) upon addition of recombinant proteins (identified in panel F) to the culture media. One-way analysis of variance (ANOVA) with Tukey’s post hoc analysis; *p* < 0.05, p < 0.01, ns: not significant. Data are presented as mean ± SEM; *n* = 3 independent experiments, each with three wells per condition. Each data point represents the mean of three wells per experiment. **I.** Confocal micrographs showing proliferating H3K27M_+_ DMG cells *in vitro*. H3K27M (green), EdU (red). Scale bars = 50 µm.

Previously identified activity-regulated, growth-promoting neuron-to-glioma signaling factors in forebrain and retinal explants include neuroligin-3 (NLGN3)(*3*, *4, 11*) and brain-derived neurotrophic factor (BDNF)( *3*, *6, 11*). Filtering hindbrain CM by molecular size revealed a growth-promoting effect of factors in the 10–100kDa range, suggesting the growth-promoting factors in this assay represent macromolecules such as proteins. Testing the role of known activity-regulated glioma mitogens NLGN3 and BDNF, we found that neither sequestering NLGN3 with neurexin-1(NRXN1) (*3*) nor inhibiting the BDNF receptor TrkB with ANA-12 fully reverted the effects of serotonergic neuronal activity-regulated secreted factors on glioma proliferation to baseline levels (***fig. S3G-H***), suggesting the involvement of additional activity-dependent secreted factors. Liquid chromatography–mass spectrometry analysis of CM revealed a distinct profile enriched in proteins associated with synaptic plasticity and transmission after serotonergic neuronal stimulation (***Fig. 3D***, ***fig. S3I-J***). Further filtering identified 13 soluble factors (***Fig. 3E-F***), some of which overlapped with previously studied factors secreted in response to glutamatergic neuronal stimulation (*3*). We focused on a subset not previously identified or tested: ICAM-1, NCAM-1, SEMA4B, and CLSTN1. Proliferation assays utilizing recombinant proteins revealed that among these, NCAM-1 and ICAM-1 robustly increased glioma cell proliferation in two distinct patient-derived glioma models, while SEMA4B, and CLSTN1 had no effect on glioma proliferation (***Fig. 3G-I***). These findings reveal a previously unrecognized set of paracrine factors released in the brainstem in response to activity of serotonergic neurons that drive glioma proliferation, adding to the list of activity-regulated paracrine factors promoting glioma growth.

### 5HT2A Activation Mediates the Glioma Response to Serotonergic Projections

We next explored the effects of serotonin on glioma proliferation. First, we treated patient-derived IDH-wildtype glioblastoma and H3K27M-mutant DMG patient-derived cell cultures with serotonin over a range of concentrations and found a dose-dependent increase in glioma proliferation rates (***Fig. 4A-C***, ***fig. S4A-B***), consistent with an early report in a glioblastoma model(*22*).

**Fig. 4:**
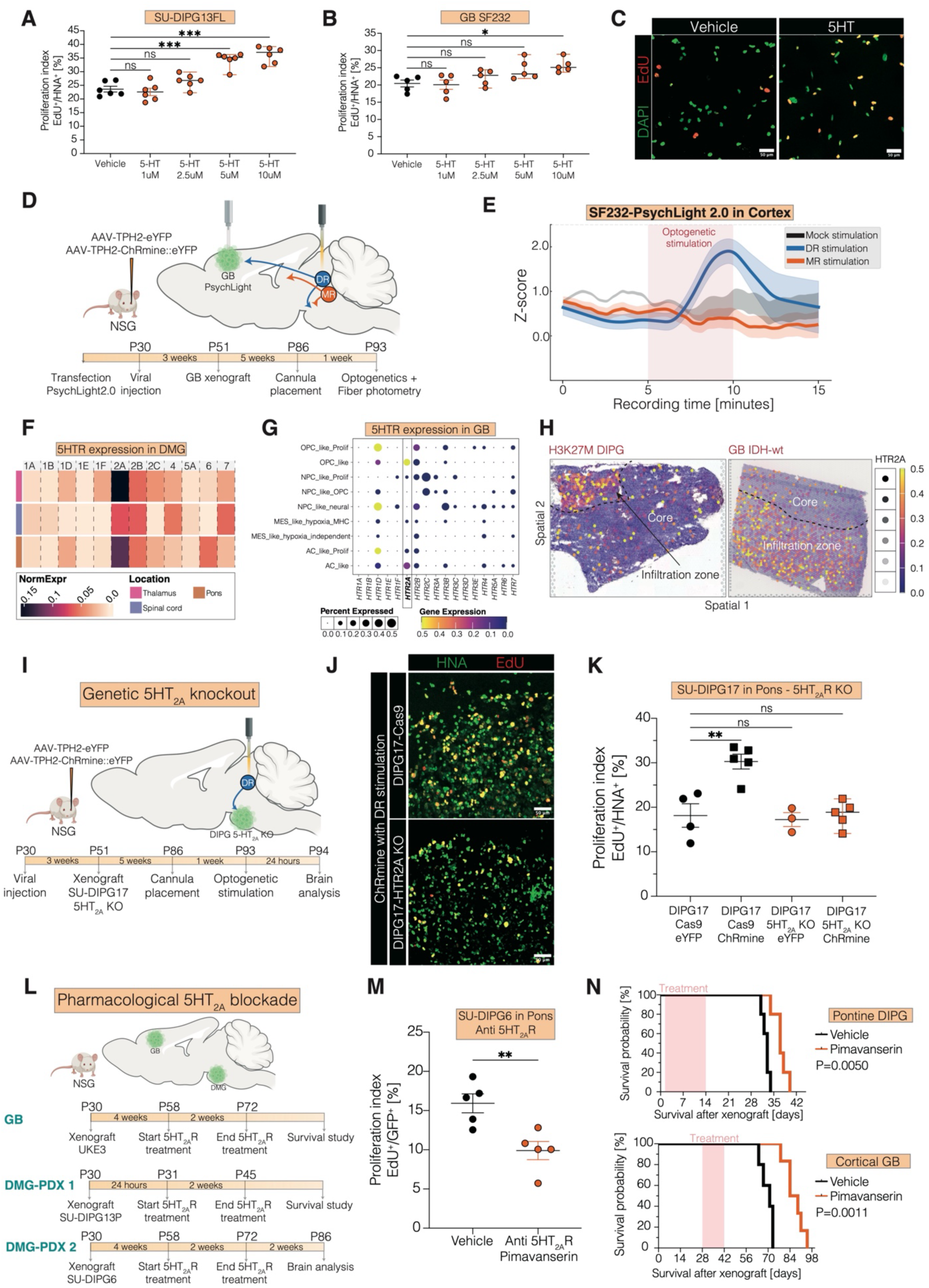
5HT_2A_ receptor activation mediates glioma response to 5HT neuronal activity. **A.** Proliferation index (EdU⁺/DAPI⁺) of monocultures of a patient-derived DMG line treated with varying doses of 5HT. One-way ANOVA with Tukey’s post hoc test; ***p < 0.001, **p < 0.01, *p < 0.05, ns: non-significant. Data are presented as mean ± SEM. n=five to six independent experiments, each with three wells per condition; each data point represents the mean of three wells per condition for a given experiment. **B.** Proliferation index (EdU⁺/DAPI⁺) of monocultures of a patient-derived adult glioblastoma line treated with varying doses of 5HT. One-way ANOVA with Tukey’s post hoc test; ***p < 0.001, **p < 0.01, *p < 0.05, ns: non-significant. Data are presented as mean ± SEM. n=five to six independent experiments, each with three wells per condition; each data point represents the mean of three wells per condition for a given experiment. **C.** Confocal micrographs showing proliferating glioblastoma cells *in vitro*. DAPI (blue), EdU (red). Scale bars = 50 µm. **D.** Schematic of the experimental paradigm for fiber photometry recordings of PsychLight2.0 fluorescence in cortical glioblastoma cells. Human glioblastoma cells (SF232) were transduced *in vitro* with PsychLight2.0 and used for xenografting. Four-week-old NSG mice (P28–30) received retro-orbital injections of rAAVPHP.eB-Tph2::eYFP or rAAVPHP.eB-Tph2::ChRmine-eYFP, followed three weeks later by xenografting of SF232-PsychLight2.0 cells into the M2 cortex. Optical ferrules were implanted into the dorsal raphe (DR) and the cortical tumor site. Fiber photometry recordings, combined with optogenetic stimulation, were performed six weeks after glioblastoma engraftment. **E.** Fiber photometry recordings showing averaged PsychLight2.0 fluorescence in M2 cortical glioblastoma cells with simultaneous mock (n=4 mice), DR (n=6 mice), and MR (n=4 mice) stimulation. **F.** Heatmap of normalized serotonergic receptor gene expression in a single cell sequencing dataset from human DMG samples, stratified by anatomical location (pons, thalamus, and spinal cord). **G.** Dot plot showing serotonergic receptor gene expression in human glioblastoma samples, depicting the proportion of glioma cells expressing each gene and the average expression level. **H.** Spatial transcriptomic data from human H3K27M and glioblastoma patient samples showing HTR2A expression patterns. **I.** Schematic of the experimental paradigm for xenografting 5HT_2A_-knockout DMG cells (SU-DIPG17-5HT_2A_ KO) with optogenetic stimulation of the DR in immunodeficient mice. Four-week-old NSG mice (P28–30) were retro-orbitally injected with rAAVPHP.eB-Tph2::eYFP or rAAVPHP.eB-Tph2::ChRmine-eYFP, followed three weeks later by xenografting of patient-derived SU-DIPG17-5HT_2A_ KO cells into the ventral pons. After five weeks of tumor growth, optical ferrules were implanted into the DR, and optogenetic stimulation was performed one week later. Mice were perfused 24 hours after stimulation. **J.** Confocal micrographs showing proliferating HNA_+_ DMG cells in the ventral pons of DR-stimulated mice xenografted with either wildtype DIPG17 (‘DIPG17-Cas9’) or 5HT_2A_ knockout cells (‘DIPG17-5HT_2A_ KO’). HNA: green, EdU: red. Scale bars = 50 µm. **K.** Proliferation index (EdU_+_/HNA_+_) of ventral pontine xenografts in mice either stimulated in DR (“ChRmine”) or mock-stimulated (“eYFP”) with wildtype DIPG (‘DIPG17-Cas9’) or 5HT_2A_ knockout cells (‘DIPG17-5HT_2A_ KO’) (n=3-5 mice/group). One-way analysis of variance (ANOVA) with Tukey’s post hoc analysis; **p < 0.01, ns: non-significant. Data=mean ± SEM. **L.** Schematic of the experimental paradigm for xenografting with 5HT_2A_ antagonist treatment (pimavanserin). Four-week-old NSG mice (P28–30) were xenografted either into the ventral pons with patient-derived DMG cells (SU-DIPG6 or SU-DIPG13P) or into the M2 cortex with patient-derived glioblastoma cells (UKE3). Chronic 5HT_2A_ antagonist treatment (i.p. 10 mg/kg) was initiated at time points specific to each cell line. **M.** Proliferation index (EdU⁺/GFP⁺) of ventral pontine xenografts in mice treated with the 5HT_2A_ antagonist pimavanserin or vehicle control (*n* = 5 mice/group). Unpaired two-tailed Welch’s *t*-test; **p < 0.01. Data are presented as mean ± SEM. **N.** Survival curves of ventral pontine xenografts (SU-DIPG13P, top) and M2 cortical xenografts (UKE3, bottom) in mice treated with either the 5HT_2A_ antagonist pimavanserin or vehicle control. Log-rank test.

To test for serotonin receptor activation in glioma *in vivo* following serotonergic neuronal activity, we packaged PsychLight2.0, a fluorescent sensor based on the structure of the 5HT_2A_ receptor to detect serotonin receptor activation, into a lentiviral backbone and transduced patient-derived glioblastoma cells. PsychLight2.0–expressing glioma cells were xenografted into cortex for fiber photometry recordings, paired with simultaneous optogenetic stimulation of dorsal raphe serotonergic neurons (***Fig. 4D***). Analysis revealed that dorsal raphe serotonergic neuronal stimulation increased PsychLight2.0 fluorescence in cortical glioma cells (***Fig. 4E***), indicating serotonin binding on glioma cells following serotonergic neuronal activity.

Next, we examined serotonin receptor expression in transcriptomic data from primary patient glioma samples. Analysis of single-cell RNA sequencing (scRNA-seq) datasets from primary DMG patient samples revealed that HTR2A is the most highly expressed serotonergic receptor gene across DMG arising in multiple anatomical locations (***Fig. 4F***, ***fig. S4C-D***) and is associated with OPC-like and AC-like malignant cell states (***fig. S4E***). Similarly, primary glioblastoma patient samples exhibited high HTR2A expression, predominantly clustering with malignant OPC- and AC-like states (***Fig. 4G***). Spatial transcriptomic data from primary patient samples further highlighted HTR2A enrichment in the glioma infiltration zone (***Fig. 4H***, ***fig. S4F***).

To test the necessity of the 5HT_2A_ receptor in the growth-promoting effects of serotonergic neuronal activity, we performed CRISPR-mediated knockout of 5HT_2A_ in patient-derived DMG cells, and then xenografted these 5HT_2A_-knockout DMG cells or *cas9*-expressing control DMG cells to the ventral pons (***Fig. 4I***). Optogenetic stimulation of dorsal raphe serotonergic neurons increased the proliferation of *cas9*-expressing control ventral pontine DMG xenografts as expected, but had no effect on the proliferation rate of 5HT_2A_-knockout ventral pontine DMG xenografts. (***Fig. 4J-K***), indicating that serotonergic signaling through the 5HT_2A_ receptor accounts completely for the growth-promoting effects of serotonergic axonal projections.

To advance clinical translation, we next tested the effects of an FDA-approved 5HT_2A_ inverse agonist (pimavanserin) on glioma proliferation rates and overall survival in models of DMG in the ventral pons or GBM in the cortex without optogenetic stimulation (***Fig. 4L***). We found reduced glioma cell proliferation rate (***Fig. 4M***) and prolonged survival in mice bearing either cortical glioblastoma or ventral pontine DMG following treatment with pimavanserin (***Fig. 4N***). Taken together, these findings identify 5HT_2A_ as the key receptor mediating serotonergic neuronal activity-regulated effects on glioma cells at sites of serotonergic projections and highlight 5HT_2A_ as a promising therapeutic target.

### Psilocybin Promotes Glioma Growth Through Direct 5HT_2A_ Activation

We next tested the effects of psilocybin, a serotonergic psychedelic with well-described functional activity at the 5HT_2A_ receptor, known to increase neuroplasticity (*23*, *24*) and currently used as an adjunctive therapy to improve quality of life in cancer patients, as well as in the treatment of major depressive disorder(*25–27*). In four distinct preclinical models of glioma, we found that a single dose of psilocybin significantly increased malignant cell proliferation in both cortical glioblastoma and ventral pontine DMG xenografts and allografts, with elevated proliferation rates persisting for up to at least two weeks post-injection (***Fig. 5A-D***, ***fig. S5A-C***).

**Fig. 5:**
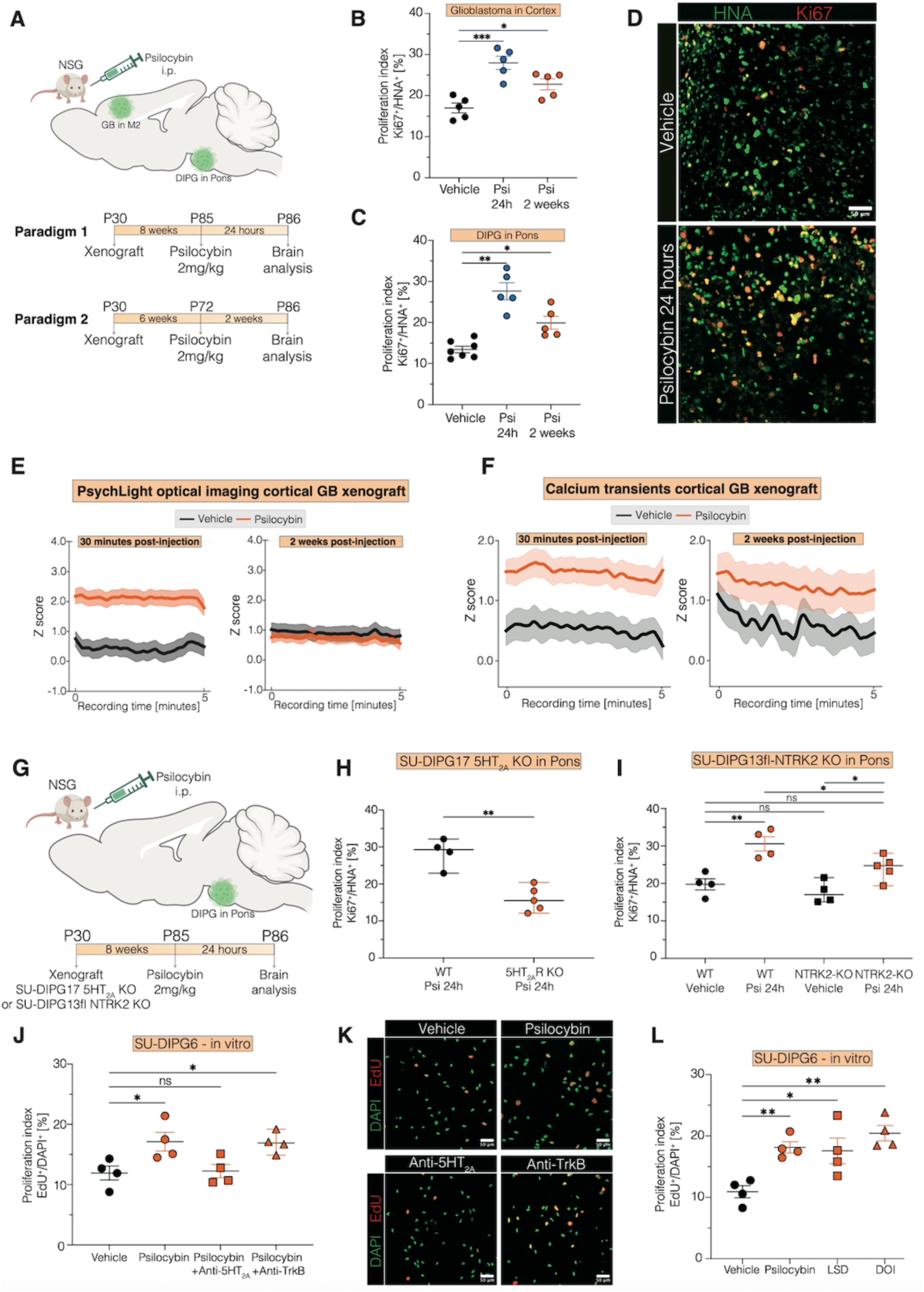
Psilocybin promotes glioma growth. **A.** Schematic of the experimental paradigm for psilocybin treatment in glioma xenograft models. Four-week-old NSG mice (P28-30) were xenografted into the M2 cortex with patient-derived glioblastoma cells or into the ventral pons with patient-derived DMG cells. Mice received intraperitoneal injections of psilocybin (2 mg/kg body weight) at 8 weeks post-xenograft (cohort 1) or 6 weeks post-xenograft (cohort 2) and were perfused either 24 hours later (cohort 1) or 2 weeks later (cohort 2). **B.** Proliferation index (Ki67⁺/HNA⁺) of M2 cortical xenografts from patient-derived glioblastoma cells (SF232) in mice treated with psilocybin (‘Psi’) or vehicle control (‘Vehicle’) (*n* = 5 mice/group). One-way analysis of variance (ANOVA) with Tukey’s post hoc test; ***p < 0.001, *p < 0.05. Data are presented as mean ± SEM. **C.** Proliferation index (Ki67⁺/HNA⁺) of ventral pontine xenografts from patient-derived DMG cells (SU-DIPG17) in mice treated with psilocybin (‘Psi’) or vehicle control (‘Vehicle’) (‘Vehicle’, *n* = 7 mice; ‘Psi 24h’ and ‘Psi 2 weeks’, *n* = 5 mice/group). One-way ANOVA with Tukey’s post hoc test; **p < 0.01, *p < 0.05. Data are presented as mean ± SEM. **D.** Confocal micrographs showing proliferating HNA_+_ DMG cells in ventral pons in vehicle-treated (upper image) and psilocybin-treated (bottom image) mice. HNA: green, Ki67: red, scale bars = 50µm. **E.** Fiber photometry recordings showing averaged PsychLight2.0 fluorescence in cortical glioblastoma cells (SF232-PsychLight2.0) at 30 minutes and 2 weeks after injection of either psilocybin or vehicle. **F.** Fiber photometry recordings showing averaged GCaMP6s-labeled calcium transients in cortical glioblastoma cells (GBM39-GCaMP6s) at 30 minutes and 2 weeks after injection of either psilocybin or vehicle. **G.** Schematic of the experimental paradigm for psilocybin treatment in glioma xenograft models. Four-week-old NSG mice (P28-30) were xenografted into the M2 cortex with patient-derived glioblastoma cells or into the ventral pons with patient-derived DMG cells. Mice received intraperitoneal injections of psilocybin (2 mg/kg body weight) at 8 weeks post-xenograft (cohort 1) or 6 weeks post-xenograft (cohort 2), and were perfused either 24 hours later (cohort 1) or 2 weeks later (cohort 2). **H.** Proliferation index (Ki67⁺/HNA⁺) of ventral pontine xenografts from patient-derived DMG cells either wildtype (SU-DIPG17 WT) or 5HT_2A_ knockout (SU-DIPG17-5HT_2A_ KO) in mice treated with psilocybin (‘Psi’) (‘WT’, n = 4 mice; ‘5HT_2A_ KO’, n = 5 mice). Unpaired two-tailed Welch’s *t*-test; **p < 0.01. Data are presented as mean ± SEM. **I.** Proliferation index (Ki67⁺/HNA⁺) of ventral pontine xenografts from patient-derived DMG cells either wildtype (SU-DIPG13fl WT) or NTRK2 knockout (SU-DIPG13fl-NTRK2 KO) in mice treated with psilocybin (‘Psi’) or vehicle control (‘vehicle’) (‘WT vehicle’, ‘WT Psi 24h’, and ‘NTRK2-KO vehicle’, n = 4 mice; ‘NTRK2-KO psi 24h’, n = 5 mice). One-way analysis of variance (ANOVA) with Tukey’s post hoc test; *p<0.05, **p < 0.01, ns: non-significant. Data are presented as mean ± SEM. **J.** Quantification of DMG cell proliferation (EdU⁺/DAPI⁺) following psilocybin treatment with or without co-administration of a 5HT2A receptor antagonist (pimavanserin) or a TrkB antagonist (ANA-12). One-way analysis of variance (ANOVA) with Tukey’s post hoc analysis; *p < 0.05, ns: non-significant. Data=mean ± SEM; n=four independent experiments, each with three wells per condition; each data point represents the mean of three wells per condition for a given experiment. **K.** Representative confocal micrographs showing glioma cell proliferation following treatment with vehicle control, psilocybin, 5HT_2A_ receptor antagonist (pimavanserin), or TrkB antagonist (ANA-12). Ki67: red, GFP: green, scale bar = 50 µm. **L.** Quantification of DMG cell proliferation (EdU⁺/DAPI⁺) following treatment with psychedelics compared to vehicle control. One-way analysis of variance (ANOVA) with Tukey’s post hoc analysis; **p < 0.01, *p < 0.05. Data=mean ± SEM; n=four independent experiments, each with three wells per condition; each data point represents the mean of three wells per condition for a given experiment.

To assess the pharmacokinetic time course of psilocybin effects on serotonergic receptor activity in glioma, we used fiber photometry to visualize fluorescence from glioma cells expressing of PsychLight2.0 (***Fig. 5E***). As expected, PsychLight2.0 *in vivo* imaging showed increased signal by 30 minutes after psilocybin administration, consistent with activation of serotonin receptors by the drug, and normalization of glioma 5HT_2A_ receptor activity to non-drug-exposed levels by two weeks after psilocybin exposure, consistent with the expectation that psilocybin should be cleared by this timepoint. However, fiber photometry to assess glioma calcium transients in GCaMP-expressing glioma cells revealed robustly increased calcium transients in glioma cells as early as 30 minutes following psilocybin administration that persisted for at least two weeks following a single psilocybin exposure (***Fig. 5F***), indicating a lasting functional effect of psilocybin on glioma.

To delineate receptor-specific contributions to psilocybin-induced glioma proliferation, we used DMG cell lines with CRISPR-mediated knockout of either 5HT_2A_ or NTRK2 (TrkB), another known psilocybin target(*28*) and a key driver of glioma proliferation(*6*). Knockout of 5HT_2A_ in glioma cells nearly abolished psilocybin-induced proliferation, whereas NTRK2 knockout partially attenuated this effect (***Fig. 5G-I***). Single-cell transcriptomic analysis of human DMG samples revealed co-expression of 5HT_2A_ and NTRK2 within individual DMG cells, primarily within malignant AC-like, OPC-like, and cycling cell populations (***fig. S5A***). In primary patient glioblastoma samples, spatial transcriptomic analyses revealed that NTRK2 expression was enriched in the OPC-like and NPC-like compartments, with highest levels in the glioma infiltration zone (***fig. S5B–D***), and co-localized with 5HT_2A_ expression (***fig. S5E***).

While blockade of the 5HT_2A_ receptor fully abrogated psilocybin-induced proliferation, pharmacological inhibition of TrkB signaling had no effect in monoculture (***Fig. 5J-K***), indicating that the NTRK2-dependent effects observed *in vivo* depend on factors in the tumor microenvironment. Further treatment of glioma monocultures revealed that, in addition to psilocybin, other serotonergic psychedelic drugs such as LSD and DOI similarly increased glioma cell proliferation (***Fig. 5L***).

Collectively, these findings demonstrate that psilocybin drives sustained glioma proliferation primarily via 5HT_2A_ receptor activation, with TrkB signaling serving as a microenvironment-dependent modulator. These results further highlight the mechanistic importance of the 5HT_2A_ receptor in glioma pathophysiology and argues for cautious clinical decision-making regarding the use of serotonergic psychedelic drugs in patients with gliomas.

### Glioma Induces Activity Changes in the Raphe Nucleus

Glioma causes a hyperexcitable microenvironment that promotes glioma-associated seizures, impairs brain functions, and further augments pathogenic neuronal inputs to the tumor(*5*, *17–19*, *29*, *30*). Recent findings demonstrate that glioma cells receive inputs from diverse neuronal types and brain regions(*15*, *16*, *31*) and reciprocally exert profound, sometimes even long-range, effects on neural activity, including distant modulation of midbrain cholinergic neurons(*21*).

Here, we investigated whether glioma cells similarly influence activity in dorsal and median raphe neurons. To label active raphe neurons in the context of glioma, we utilized TRAP2 x Ai14 mice(*32*) allografted with glioma cells into the ventral pons, administering 4-OHT four weeks post tumor cell implantation and perfusing six hours later (***Fig. 6A***). Glioma-bearing mice exhibited a significant increase in tdTomato-labeled neurons within the raphe nucleus compared to non-tumor-bearing controls, indicating heightened raphe nucleus activity in the context of glioma (***Fig. 6B-C***).

**Fig. 6:**
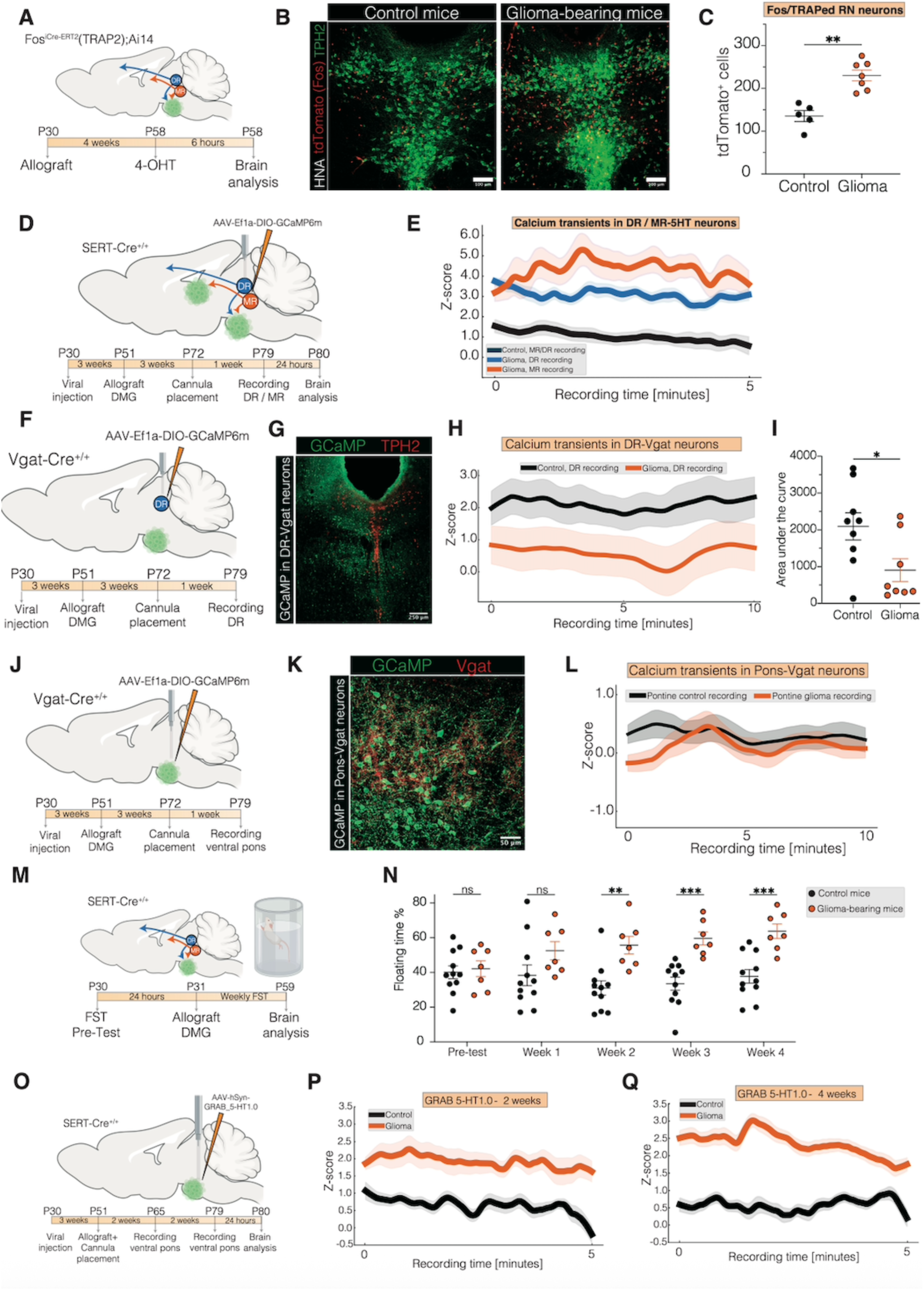
Glioma induces activity changes in the raphe nucleus and modulates depressive-like behavior. **A.** Schematic of the experimental paradigm to assess glioma-dependent activity changes in the RN. TRAP2 × Ai14 mice (P28–30) were allografted in the ventral pons with H3K27M DMG cells (‘Glioma’) or vehicle (‘Control’). Four weeks later, mice received a single intraperitoneal injection of 4-OHT (10 mg/kg) and were perfused 6 hours post-injection. **B.** Representative confocal images of the RN in control-injected (left) and DMG-bearing mice (right). TPH2: green, tdTomato: red, scale bars = 100 µm. **C.** Quantification of tdTomato⁺ cells in the RN of control versus glioma-bearing mice. Unpaired two-tailed Welch’s t-test; **p < 0.01. Data are presented as mean ± SEM. **D.** Schematic of experimental setup for *in vivo* calcium imaging of DR^5HT^ and MR^5HT^ neurons. SERT-Cre_+/+_ mice (P28–30) were injected with AAV-DJ-Ef1α-DIO-GCaMP6m into the DR or MR, and allografted with H3K27M DMG cells or vehicle three weeks later. Fiberoptic cannulae were implanted into the DR or MR one week prior to recording. **E.** Averaged GCaMP6m fiber photometry traces of DR^5HT^ and MR^5HT^ neurons in glioma-bearing and control mice at 4 weeks post-allograft (‘Control’, n = 6 mice; ‘Glioma DR’ and ‘Glioma MR’, n = 7 mice). **F.** Schematic of experimental paradigm for calcium imaging of GABAergic neurons in the DR. Vgat-Cre_+/+_ mice (P28–30) were injected with AAV-DJ-Ef1α-DIO-GCaMP6m in the DR and allografted with H3K27M DMG cells or vehicle three weeks later. Cannulae were implanted after three weeks, and recordings were performed one week later. **G.** Representative confocal image showing GCaMP6m expression in the DR of Vgat-Cre mice. GCaMP: green, TPH2: red, scale bar = 250 µm. **H.** Averaged GCaMP6m photometry traces of DR_Vgat_ neurons in glioma-bearing and control mice at 4 weeks post-allograft (‘Control’, n = 4 mice; ‘Glioma DR’, n = 6 mice). **I.** Area under the curve of photometry recordings from DR_Vgat_ neurons. Unpaired two-tailed Welch’s t-test; *p < 0.05. Data are presented as mean ± SEM. **J.** Schematic of experimental paradigm for calcium imaging of GABAergic neurons in the ventral pontine tumor microenvironment. Vgat-Cre_+/+_ mice (P28–30) were injected with AAV-DJ-Ef1α-DIO-GCaMP6m in the ventral pons and allografted with H3K27M DMG cells or vehicle three weeks later. Cannulae were implanted after three weeks, and recordings were performed one week later. **K.** Representative confocal image showing GCaMP6m expression in the ventral pons of Vgat-Cre mice. GCaMP: green, Vgat: red, scale bar = 50 µm. **L.** Averaged GCaMP6m photometry traces of ventral pontine Vgat_+_ neurons in glioma-bearing and control mice at 4 weeks post-allograft (‘Control’, n = 4 mice; ‘Glioma DR’, n = 5 mice). **M.** Schematic of forced swim test (FST) paradigm. SERT-Cre_+/+_ mice (P28–30) underwent baseline testing and were injected with controls or DMG cells into the ventral pons. Weekly FST sessions were conducted starting one-week post-allograft. **N.** Weekly analysis of immobility in control- and glioma-bearing mice. Unpaired two-tailed Welch’s t-test; **p < 0.01, ***p < 0.001, ns: non-significant. Data are presented as mean ± SEM. **O.** Schematic of the experimental setup for *in vivo* fluorescence imaging of 5HT release within the glioma microenvironment. SERT-Cre_+/+_ mice (P28–30) were injected with AAV-hSyn- GRAB_5-HT1.0 into ventral pons, and allografted with H3K27M DMG cells or vehicle three weeks later. Fiberoptic cannulae were implanted at the time of glioma cell allografting. **P.** Averaged GRAB_5-HT1.0 fiber photometry traces in glioma-bearing and control mice at 2 weeks post-allograft (‘Control’, n = 5 mice; ‘Glioma’, n = 6 mice). **Q.** Averaged GRAB_5-HT1.0 fiber photometry traces in glioma-bearing and control mice at 4 weeks post-allograft (‘Control’, n = 5 mice; ‘Glioma’, n = 6 mice).

To further investigate raphe neuronal activity changes, we performed fiber photometry of neuronal calcium transients of dorsal and median raphe serotonergic neurons expressing the genetically encoded calcium indicator GCaMP6m (***Fig. 6D***). Four weeks after ventral pontine or thalamic allograft, serotonergic neurons in both raphe nuclei exhibited increased calcium transients relative to non-tumor-bearing controls (***Fig. 6E***).

Given the transcriptomic, spatial, functional, and subtype diversity of neurons in the raphe nuclei (*20*, *33*, *34*), we sought to obtain a more granular view about the possibly heterogeneous response of raphe neurons to distant glioma cells. One established mechanism of glioblastoma-induced elevations in cortical glutamatergic neuronal activity is decreased inhibitory inputs from GABAergic inhibitory neurons(*35*). We therefore hypothesized that altered GABAergic inhibitory neuronal function may contribute to the elevated serotonergic neuronal activity in the raphe. We assessed GABAergic neuronal activity in the raphe nuclei using fiber photometry to monitor calcium transients of Vgat_+_ GABAergic neurons in mice bearing DMG in the ventral pons or in non-tumor-bearing control mice (***Fig. 6F-G***). In mice with ventral pontine DMG tumors, we discovered a marked reduction in calcium transients in Vgat+ neurons in the dorsal raphe at four weeks post-glioma cell implantation, indicating reduced inhibitory neuronal activity in the raphe (***Fig. 6H-I***) which could be due to either reduced activity or reduced numbers of GABAergic inhibitory neurons.

In glioblastoma, GABAergic neurons are lost from the tumor microenvironment(*35*), but this does not occur in DMG(*7*), consistent with the demonstrated growth-promoting role of GABAergic neurons in DMG specifically (*7*). Given the above finding of reduced GABAergic neuronal activity in the raphe when DMG is present in the ventral pons, we next assessed the activity of GABAergic neurons in the ventral pontine DMG microenvironment. Fiber photometry recordings from the ventral pontine DMG microenvironment revealed no differences between glioma-bearing and control-injected animals (***Fig. 6J-L***), indicating no reduction of GABAergic neuronal activity in the ventral pons, as expected(*7*). Taken together, these results underscore region-specific effects of ventral pontine DMG on GABAergic neuronal activity, with reduction in the dysregulated raphe nuclei but preservation of DMG growth-promoting GABAergic activity in the local tumor microenvironment of the ventral pons(*7*).

Given this observed dysregulation of raphe neuronal activity in the context of glioma, and the high rate of depression and other psychiatric symptoms that patients with gliomas experience(*36*), we utilized mouse behavioral tests of brain functions to which the finely regulated activity of the raphe nuclei contribute, such as active coping(*37*). We used the forced swim test to assess depressive-like behavior at a time point (4 weeks post–tumor cell implantation) when the mice did not display gross motor function deficits (***Fig. 6M***). We found that in mice bearing ventral pontine DMG tumors, the observed alterations in raphe nuclei neuronal activity described above were accompanied by increased immobility/decreased active coping among glioma-bearing mice compared to non-tumor-bearing mice (***Fig. 6N***).

Exploring the translational problem of depression in the setting of a tumor for which serotonergic signaling promotes cancer growth, we tested the effects of mirtazapine, a tetracyclic antidepressant with 5HT_2A_ antagonist properties. Mice bearing ventral pontine DMG xenografts exhibited reduced glioma cell proliferation rate following 4 weeks of treatment with mirtazapine (***fig. S6A-B***), underscoring its potential to reduce glioma growth and also highlighting mirtazapine as a safe choice from a tumor growth perspective for treatment of depression in the context of this cancer.

Finally, we assessed whether the glioma-associated changes in raphe nucleus neuronal activity described above were associated with elevated serotonin release in the glioma microenvironment over time. Using the GRAB_5HT1.0 sensor to visualize serotonin binding to 5HT receptors together with fiber photometry in the ventral pontine tumor microenvironment, we measured progressively increased serotonin release in the ventral pontine glioma microenvironment over the disease course (***Fig. 6O–Q***), highlighting the vicious, feed-forward cycle of growth-promoting effects of serotonin on glioma cells and glioma-induced increases in serotonergic neuronal activity resulting in progressively elevated serotonin release.

## Discussion

High-grade gliomas remain among the most lethal tumors, with glioblastoma representing the deadliest primary brain malignancy in adults and DMG in children. Tumor–microenvironment interactions have emerged as key determinants of glioma progression, with neuron-glioma interactions emerging as a major driver and hallmark of cancer(*2*, *38*). Previous work has implicated glutamatergic(*3*, *5*, *6*, *8*, *10*), GABAergic(*7*), and cholinergic neurons(*15*, *16*, *21*) in promoting glioma growth. Our findings now extend this principle to include serotonergic neurons originating in the dorsal and median raphe, underscoring the remarkable diversity and complexity of neuron–glioma interactions.

Serotonin is an abundant neuromodulator in the brain and serotonergic circuitry plays central roles in neuropsychiatric brain functions and dysfunction. However, its significance in glioma has remained largely unexplored. An early study reported that serotonin exposure promotes both proliferation and migration of glioblastoma cells(*22*), and more recent work has demonstrated that cortical glioblastomas receive serotonergic inputs from the raphe nuclei (*15*, *16*, *31*), yet the direct functional interactions between glioma cells and long-range serotonergic projections have to date been understudied.

Here, we demonstrate that dorsal and median raphe nucleus serotonergic neuronal projections drive the proliferation and progression of both glioblastoma and DMG in a circuit-dependent manner. This growth-promoting effect at serotonergic projection sites such as cortex, thalamus and ventral pons is mediated by 5HT_2A_ receptor activation on glioma cells, triggered by 5HT release by serotonergic axons. In the region of the raphe nuclei, activity-regulated paracrine factors including NCAM1 and ICAM1 also contribute to the growth-promoting effects of serotonergic neuronal activity. Pharmacological blockade of 5HT_2A_ suppresses tumor proliferation and extends survival *in vivo*, whereas psilocybin-induced activation of 5HT_2A_ elicits sustained calcium transients and persistently enhances glioma growth. Importantly, we find that serotonergic neuronal activity-driven glioma proliferation is highly circuit-dependent, indicating that brain tumor location influences the recruitment of specific neuronal projections in accordance with established neurobiological principles.

The identification of 5HT_2A_ as a pivotal mediator of 5HT activity-driven glioma growth in both glioblastoma and DMG presents a promising therapeutic target. The existence of FDA-approved, brain-penetrant 5HT_2A_ antagonists such as pimavanserin and mirtazapine highlights the possibility of repurposing these agents in glioma treatment, which should be further studied in prospective clinical trials. Conversely, our results caution against the use of 5HT_2A_ activators such as psilocybin in patients with glioma, given the profound and sustained tumor growth-promoting effects observed.

Gliomas dysregulate raphe neurons during disease progression, establishing a feed-forward loop that exacerbates tumor progression and associates with reduced active coping behavior in glioma-bearing mice. Numerous mechanisms by which gliomas induce the hyperexcitability of glutamatergic neurons have been described, including local secretion of glutamate via the system xC glutamate-cysteine exchanger(*18*, *29*) secretion of synaptogenic factors such as glypican-3(*19*, *30*) and thrombospondin-1(*17*), and dropout of inhibitory neurons in the local tumor microenvironment of glioblastoma, the mechanisms through which gliomas alter serotonergic neuronal activity – particularly at a distance – remain to be elucidated. Our findings of diminished GABAergic inhibitory neuronal activity suggest one component of mechanisms driving progressive serotonergic neuronal dysregulation. Similar mechanisms have been reported in other cancer models at even further distance, such as breast cancer causing a loss of inhibitory input to – and consequently elevated activity of – hypothalamic paraventricular nucleus neurons expressing corticotrophin releasing hormone(*39*). Long-range, brain-wide and brain-body signaling between cancer and the nervous system is thus emerging as an important dimension of cancer neuroscience. Further studies are needed to dissect the molecular pathways and structural remodeling underlying these processes, which may reveal new avenues for therapeutic intervention beyond direct tumor targeting. Furthermore, given the high rates of depression and anxiety across a wide range of cancer patient populations(*40*), it is important to study how other cancers - including cancers outside of the brain - might similarly induce raphe neuronal dysregulation. Serotonergic neuronal regulation of functions such as active coping is complex and finely regulated by distinct subpopulations of serotonergic neurons, with some subpopulations of serotonergic neurons promoting active coping and others decreasing active coping(*20*). Future work will be required to understand more about precisely which serotonergic neurons are dysregulated by gliomas and other cancers. Understanding the details of how gliomas and possibly other cancers dysregulate raphe neurons is pressing, as developing strategies to normalize raphe functions may mitigate the quality of life-impairing neuropsychiatric symptoms commonly experienced by patients with gliomas(*36*) and other cancers(*40*).

Taken together, the findings presented here establish dorsal and median raphe serotonergic neurons and their projections as key drivers of glioma progression, offering new mechanistic insight and highlighting 5HT_2A_ antagonism as a promising therapeutic strategy to disrupt neuron–glioma crosstalk and improve outcomes for these presently lethal brain cancers.

## Funding

National Institute of Neurological Disorders and Stroke R01NS092597 (MM)

NIH Director’s Pioneer Award DP1NS111132 (MM)

National Cancer Institute P50CA165962, R01CA258384, and U19CA264504 (MM)

Cancer Research UK (MM)

Cancer Grand Challenges OT2CA278688 and CGCATF-2021/100012 (MM)

Gatsby Charitable Foundation: Gatsby Initiative in Brain Development and Psychiatry (MM and KD)

Oscar’s Kids Foundation (MM)

McKenna Claire Foundation (MM)

Will Irwin Research Fund of the Pediatric Cancer Research Foundation (MM)

Yuvaan Tiwari Foundation (MM)

Austin Strong Foundation (MM)

Avery Huffman DIPG Foundation (MM)

Chadtough Defeat DIPG (MM)

Maternal and Child Health Research Institute at Stanford University Postdoctoral Award (BY),

Dean’s Postdoctoral Fellowship at Stanford University (BY)

Alex’s Lemonade Stand Foundation 1278926 (RD)

German Research Foundation 539349120 (RD) AE Foundation (KD)

National Institutes of Health grant P50DA042012 (KD)

Solving Kids’ Cancer/The Bibi Fund (MGF) Jonah Finn Foundation (MGF)

Caroline Mortimer Fund (MGF)

Career Award for Medical Scientists from the Burroughs Wellcome Fund (MGF)

Distinguished Scientist Award from the Sontag Foundation (MGF)

Alex’s Lemonade Stand Foundation ‘A’ Award (MGF)

Claudia Adams Barr Program in Innovative Cancer Research/Dana-Farber Cancer Institute (MGF)

National Institutes of Health Director’s New Innovator Award DP2NS127705 (MGF)

## Author Contributions

Conceptualization: RD and MM

Methodology: RD, BY, AbR, PJW, RoM, BDH, MGF, KD

Investigation: RD, ReM, KS, AlR, SW, YAY, LS, CAB, CLC

Visualization: RD, CAB, DHH

Funding acquisition: MM

Project administration: RD

Supervision: MM

Writing – original draft: RD and MM

## Data and material availability

All raw data presented in the manuscript are available from the corresponding author upon reasonable request. Single-cell RNA-sequencing data for the analysis of DMG patient samples were obtained from GSE184357 (SMART-seq data only, n=18 patients)(*41*). Single-cell RNA-sequencing data for the analysis of adult glioblastoma patient samples and spatial transcriptomics data were obtained from Datadryad (https://doi.org/10.5061/dryad.h70rxwdmj)(42), and the associated code is available at https://github.com/theMILOlab.

---

**Fig. S1:**
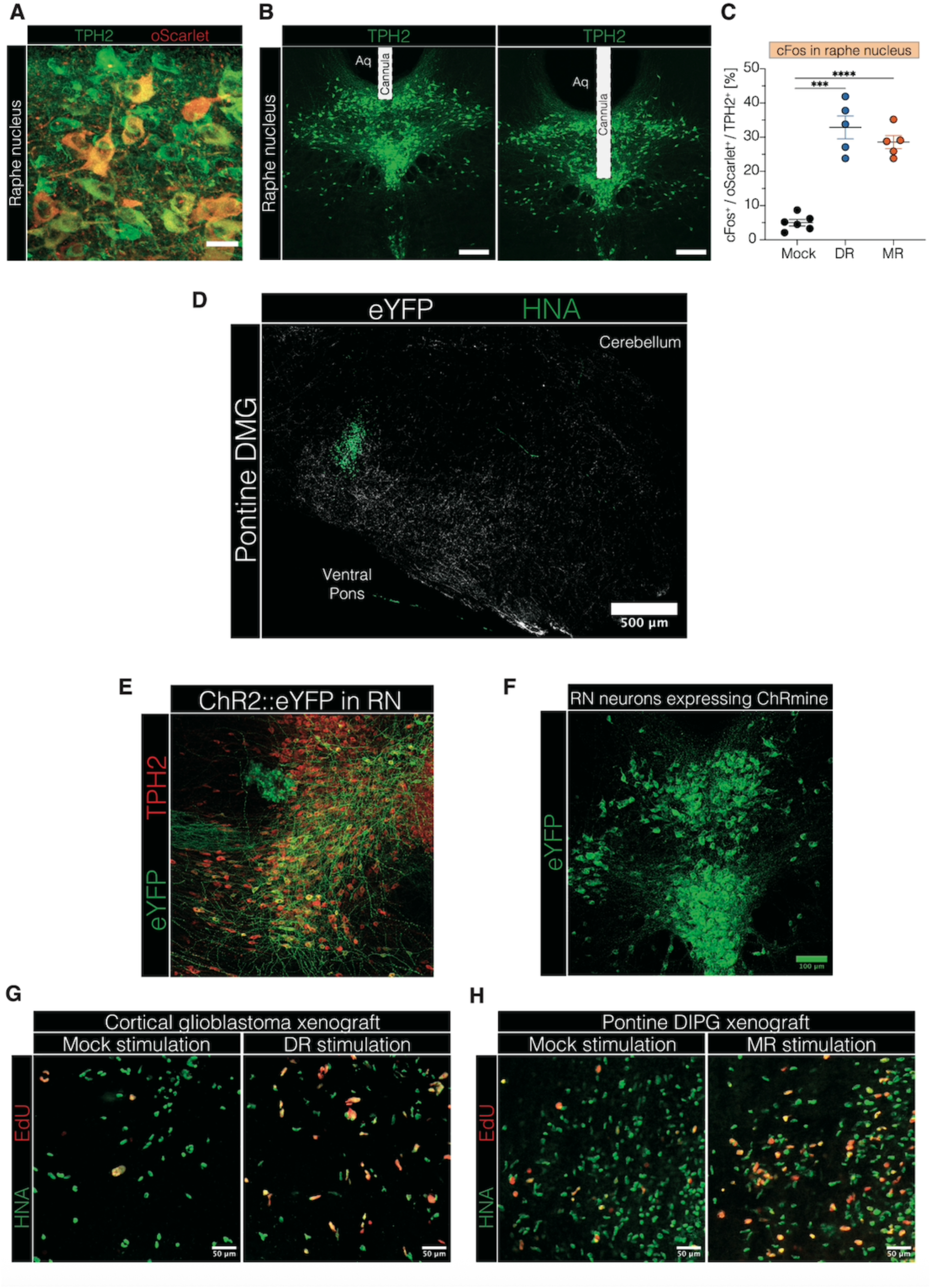
5HT neuron expression within the raphe nucleus and activity-mediated glioma proliferation effects. **A.** Representative confocal image showing oScarlet expression in the raphe nucleus of SERT-Cre x Ai230 mice. oScarlet: red, TPH2: green, scale bar = 20 µm. **B.** Confocal micrographs showing representative fiberoptic cannula placement for optogenetic experiments in the DR (left) and MR (right). TPH2: green, scale bar = 200 µm. **C.** Quantification of cFos expression following optogenetic stimulation of 5HT neurons in the DR or MR compared to mock-stimulated controls in SERT-Cre x Ai230 mice during a single stimulation session (t = 30 minutes) (‘Mock’, ‘DR’, and ‘MR’, n = 5 mice). One-way ANOVA with Tukey’s post hoc test; ****p < 0.0001, ***p < 0.001. Data are presented as mean ± SEM. **D.** Representative confocal image showing virally labeled 5HT neurons within the ventral pontine glioma microenvironment in SERT-Cre mice. eYFP: red, HNA: green, scale bar = 500 µm. **E.** Representative confocal micrograph showing viral labeling (AAV-DJ-EF1α-DIO-hChR2(H134R)::eYFP) of MR^5HT^ neurons in a sagittal section. eYFP: green, TPH2: red, DAPI: white. **F.** Representative confocal micrograph showing viral labeling (rAAVPHP.eB-Tph2::ChRmine-eYFP) of RN_5HT_ neurons. eYFP: green, scale bar = 100µm. **G.** Confocal micrographs showing proliferating HNA_+_ glioblastoma cells in M2 cortex in mock-stimulated (left image) and DR-stimulated (right image) mice. HNA: green, EdU: red, scale bars = 50µm. **H.** Confocal micrographs showing proliferating HNA_+_ DMG cells in ventral pons in mock-stimulated (left image) and MR-stimulated (right image) mice. HNA: green, EdU: red, scale bars = 50µm.

**Fig. S2:**
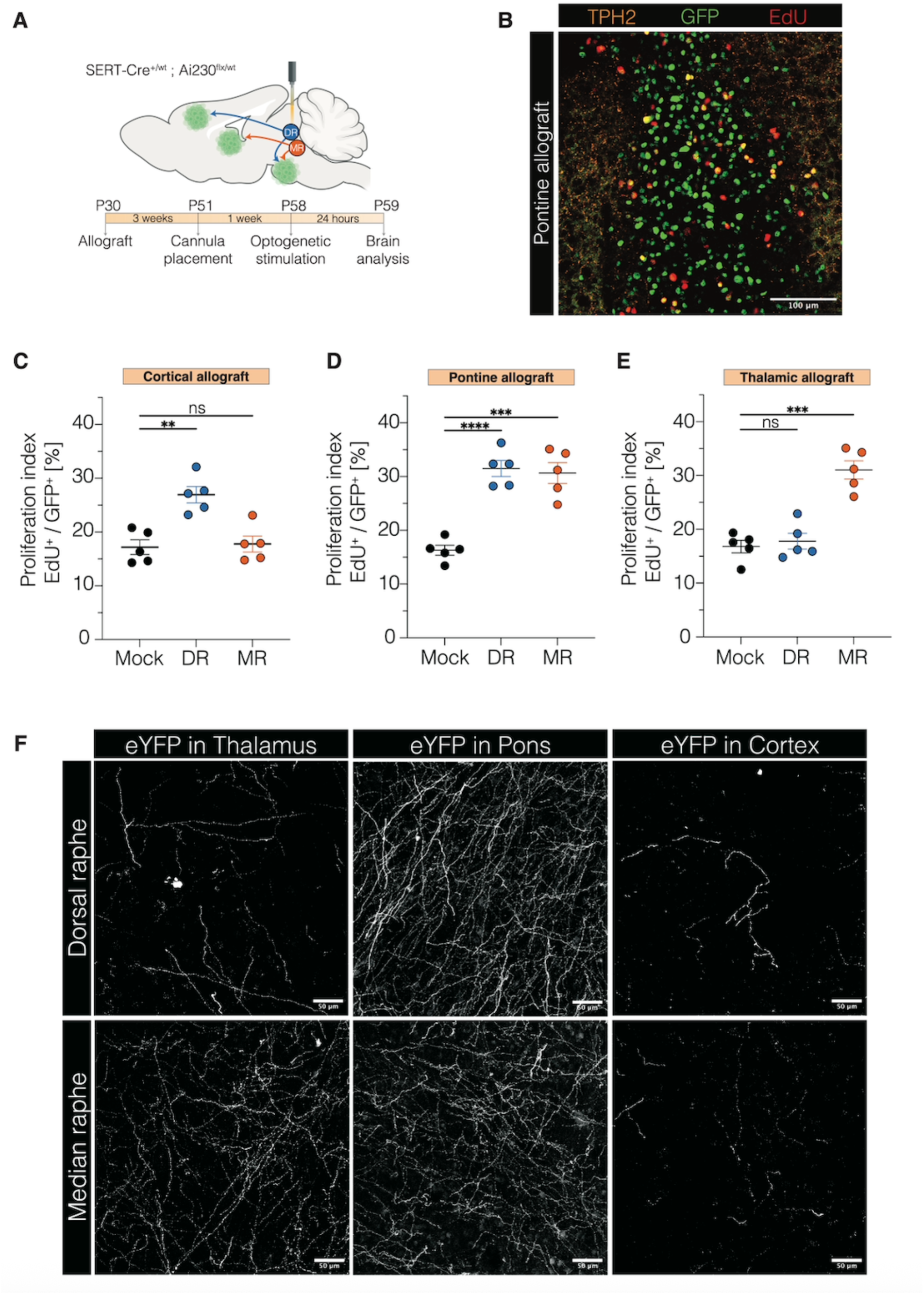
5HT neuronal activity promotes glioma proliferation in a circuit-dependent manner across distinct anatomical locations. **A.** Schematic of experimental paradigm for optogenetic stimulation of DR^5HT^ and MR^5HT^ neurons in mice bearing H3K27M DMG. Four-week-old SERT-Cre_+/wt_ x Ai230_flx/wt_ mice (P28-30) were allografted with a H3K27M MADR tumor model into neocortex (M2), thalamus, or ventral pons, with optic ferrule placement into the DR or MR three weeks after allografting. Optogenetic stimulation for 30 minutes of the DR or MR was performed four weeks after allografting, followed by perfusion 24 hours after stimulation. **B.** Representative confocal image showing GFP-labeled DMG cells in the ventral pons and 5HT projections labeled by TPH2. GFP: green, TPH2: orange, EdU: red, scale bar = 100 µm. **C.** Proliferation index (EdU_+_/GFP_+_) of cortical allografts in mice either stimulated in DR or MR or mock-stimulated (“Mock”) (Mock, DR, and MR, n=5 mice/group). Unpaired two-tailed Welch’s t-test; **p < 0.01, ns: non-significant. Data=mean ± SEM. **D.** Proliferation index (EdU_+_/GFP_+_) of ventral pontine allografts in mice either stimulated in DR or MR or mock-stimulated (“Mock”) (Mock, DR, and MR, n=5 mice/group). Unpaired two-tailed Welch’s t-test; ****p < 0.0001, ***p < 0.001. Data=mean ± SEM. **E.** Proliferation index (EdU_+_/GFP_+_) of thalamic allografts in mice either stimulated in DR or MR or mock-stimulated (“Mock”) (Mock, DR, and MR, n=5 mice/group). Unpaired two-tailed Welch’s t-test; ***p < 0.001, ns: non-significant. Data=mean ± SEM. **F.** Representative confocal images showing virally labeled 5HT projections from the MR (left images) and DR (right images) in the thalamus, ventral pons, and cortex. eYFP: white, scale bar = 50 µm.

**Fig. S3:**
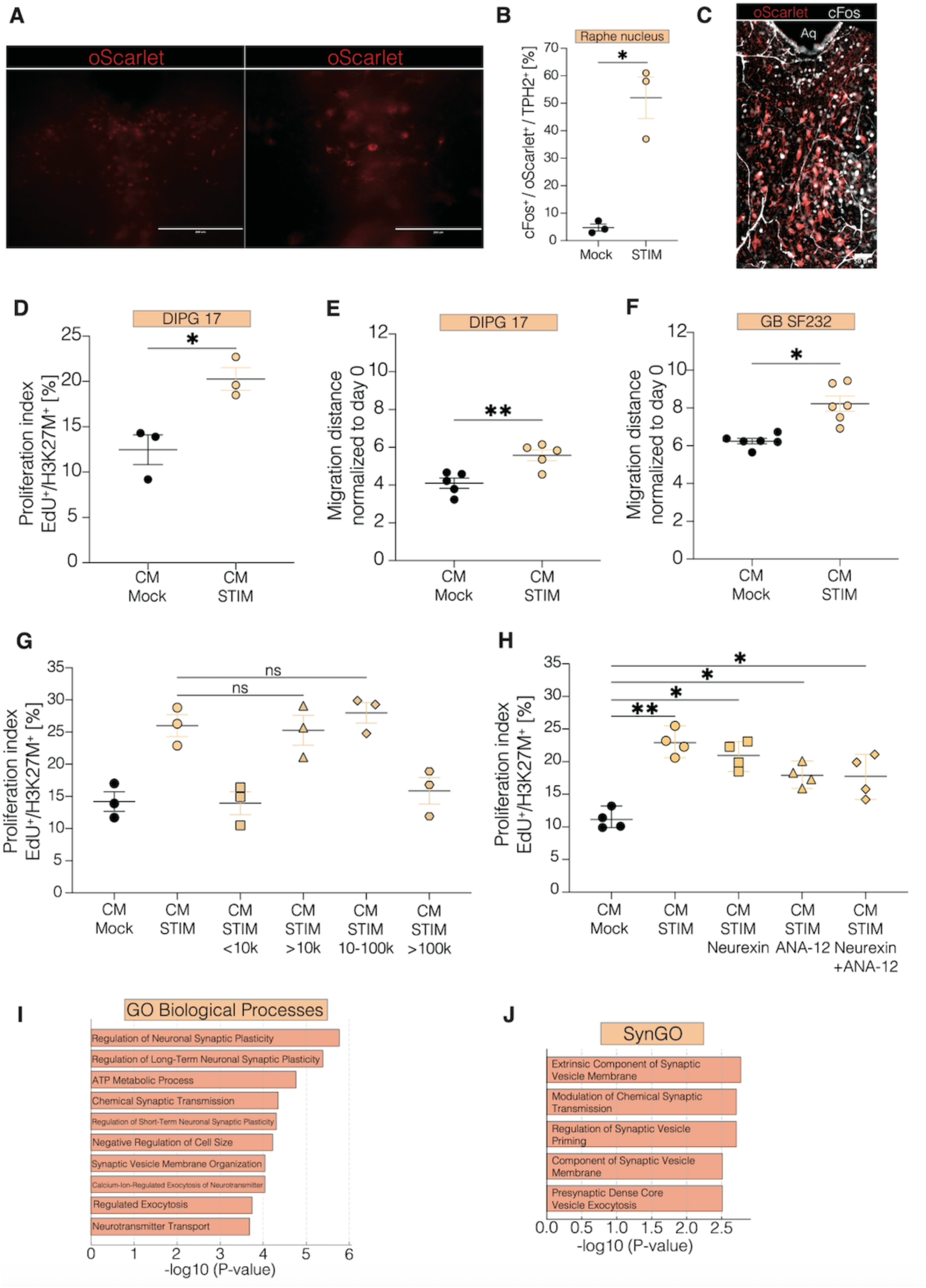
Conditioned media derived from 5HT neuronal activity in the raphe nucleus promotes glioma cell proliferation and migration. **A.** Representative confocal images showing local oScarlet-labeled ChRmine expression in the raphe nucleus in live *ex vivo* active slices. oScarlet: red, scale bar = 200 µm. **B.** Quantification of cFos expression following *ex vivo* optogenetic stimulation of 5HT neurons in the RN (“STIM”) compared to mock-stimulated controls (“Mock”) in active slices of SERT-Cre x Ai230 mice. Unpaired two-tailed Welch’s t-test; *p < 0.05. Data are presented as mean ± SEM. **C.** Confocal micrograph showing cFos expression in the raphe nucleus fixed active slices. oScarlet: red, cFos: white, scale bar = 50 µm. **D.** Quantification of DMG cell proliferation (EdU_+_/H3K27M_+_) when adding CM after *ex vivo* stimulation of RN compared to CM from mock-stimulated slices. Unpaired two-tailed Welch’s t-test; *p < 0.05. Data=mean ± SEM; n=three independent experiments, each with three wells per condition; each data point represents the mean of three wells per condition for a given experiment. **E.** Quantification of DMG cell migration after 72hours when adding CM after *ex vivo* stimulation of RN compared to CM from mock-stimulated slices. Unpaired two-tailed Welch’s t-test; **p < 0.01. Data=mean ± SEM; n=five independent experiments, each with three wells per condition; each data point represents the mean of three wells per condition for a given experiment. **F.** Quantification of glioblastoma cell migration after 72hours when adding CM after *ex vivo* stimulation of RN compared to CM from mock-stimulated slices. Unpaired two-tailed Welch’s t-test; *p < 0.05. Data=mean ± SEM; n=five independent experiments, each with three wells per condition; each data point represents the mean of three wells per condition for a given experiment. **G.** Proliferation index (EdU_+_/H3K27M_+_) of a DMG cell line (“SU-DIPG17”) after fractionation of the CM by molecular weight. One-way analysis of variance (ANOVA) with Tukey’s post hoc analysis; ns: non-significant. Data=mean ± SEM; n=three independent experiments, each with three wells per condition; each data point represents the mean of three wells per condition for a given experiment. **H.** Proliferation index (EdU_+_/H3K27M_+_) of a DMG cell line (“SU-DIPG17”) after adding a NLGN3-antagonist (Neurexin) and TrkB-inhibitor (ANA-12) to the CM. One-way analysis of variance (ANOVA) with Tukey’s post hoc analysis; **p < 0.01, *p < 0.05, ns: non-significant. Data=mean ± SEM; n=three independent experiments, each with three wells per condition; each data point represents the mean of three wells per condition for a given experiment. **I.** Gene ontology terms for biological processes identified in the CM from stimulated active slices, based on the top 50 upregulated proteins. **J.** Gene ontology terms using SynGO analysis of the CM from stimulated active slices, based on the top 50 upregulated proteins.

**Fig. S4:**
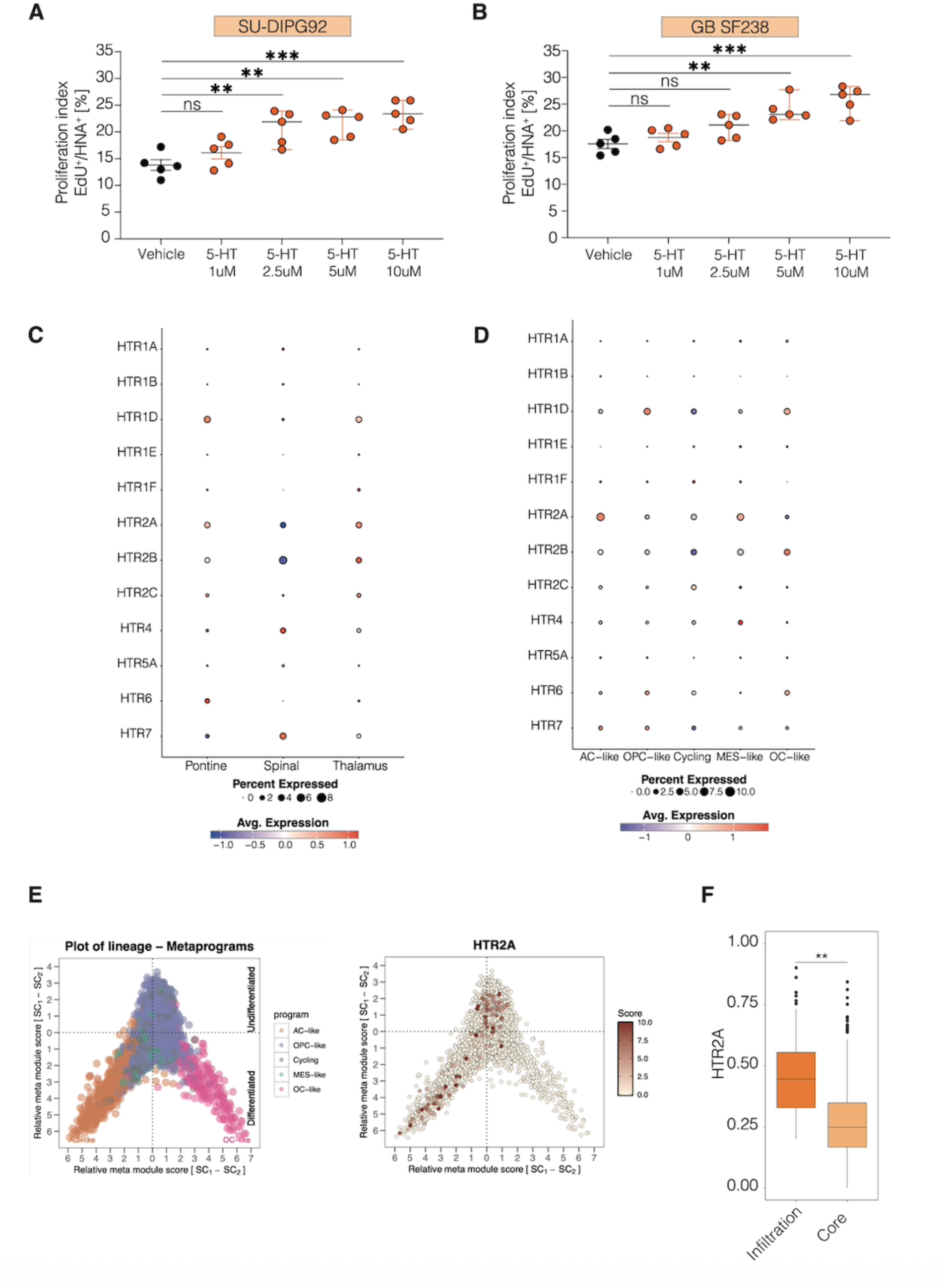
HTR2A is the most upregulated serotonergic receptor in both diffuse midline glioma and glioblastoma. **A.** Proliferation index (EdU⁺/DAPI⁺) of monocultures of a patient-derived DMG line treated with varying doses of 5HT. One-way ANOVA with Tukey’s post hoc test; ***p < 0.001, **p < 0.01, *p < 0.05, ns: non-significant. Data are presented as mean ± SEM. n=five to six independent experiments, each with three wells per condition; each data point represents the mean of three wells per condition for a given experiment. **B.** Proliferation index (EdU⁺/DAPI⁺) of monocultures of a patient-derived adult glioblastoma line treated with varying doses of 5HT. One-way ANOVA with Tukey’s post hoc test; ***p < 0.001, **p < 0.01, *p < 0.05, ns: non-significant. Data are presented as mean ± SEM. n=five to six independent experiments, each with three wells per condition; each data point represents the mean of three wells per condition for a given experiment. **C.** Illustration of serotonergic receptor gene expression in human DMG samples, showing the number of glioma cells expressing the gene and the average expression level across ventral pons, thalamus, and spinal cord. **D.** Illustration of serotonergic receptor gene expression in human DMG samples, showing the number of glioma cells expressing the gene and the average expression level across different glioma cell states. **E.** HTR2A expression level in malignant H3K27M+ malignant single cells projected on the metaprograms (x axis) and stemness (undifferentiated to differentiated; y axis) scores. **F.** Comparison of HTR2A expression between the tumor core and infiltration zone based on spatial transcriptomic data from human glioblastoma samples. Unpaired two-tailed Welch’s t-test; **\*\***p < 0.01.

**Fig. S5:**
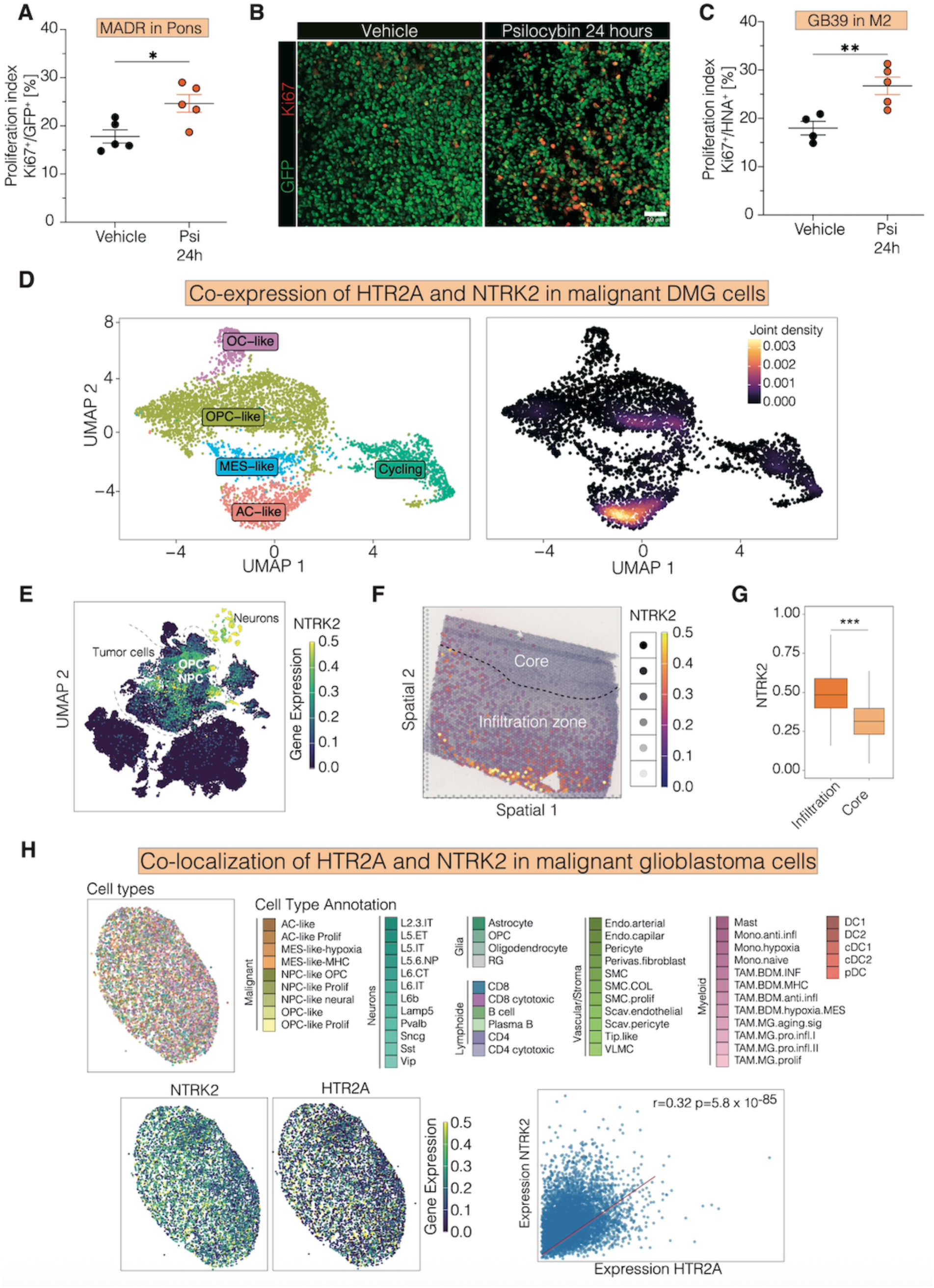
Psilocybin effects on glioma cell proliferation and co-expression of HTR2A and NTRK2. **A.** Proliferation index (Ki67⁺/GFP⁺) of ventral pontine allografts (MADR) in mice treated with psilocybin (‘Psi’) or vehicle control (‘Vehicle’) (*n* = 5 mice/group). Unpaired two-tailed Welch’s t-test; *p < 0.05. Data are presented as mean ± SEM. **B.** Representative confocal micrograph showing proliferation rates in ventral pontine allografts following treatment with psilocybin or vehicle control. Ki67: red, GFP: green, scale bar = 50 µm. **C.** Proliferation index (Ki67⁺/HNA⁺) of cortical xenografts (GB39) in mice treated with psilocybin (‘Psi’) or vehicle control (‘Vehicle’) (‘Psi’, *n* = 5 mice; ‘Vehicle’, *n* = 4 mice). Unpaired two-tailed Welch’s t-test; **p < 0.01. Data are presented as mean ± SEM. **D.** Single-cell analysis of human DMG samples showing co-expression of HTR2A and NTRK2 within the same glioma cells, clustered by malignant metaprograms. **E.** UMAP plot illustrating NTRK2 expression across different cell types and cell states based on single-cell sequencing datasets from human glioblastoma samples. **F.** Analysis of spatial NTRK2 expression in human glioblastoma samples, **G.** Expression levels of NTRK2 compared between the tumor infiltration zone and core. **H.** Spatial transcriptomic analysis of human glioblastoma samples showing cell type–dependent expression of HTR2A and NTRK2, and their spatial co-localization and correlation within the tumor microenvironment.

**Fig. S6:**
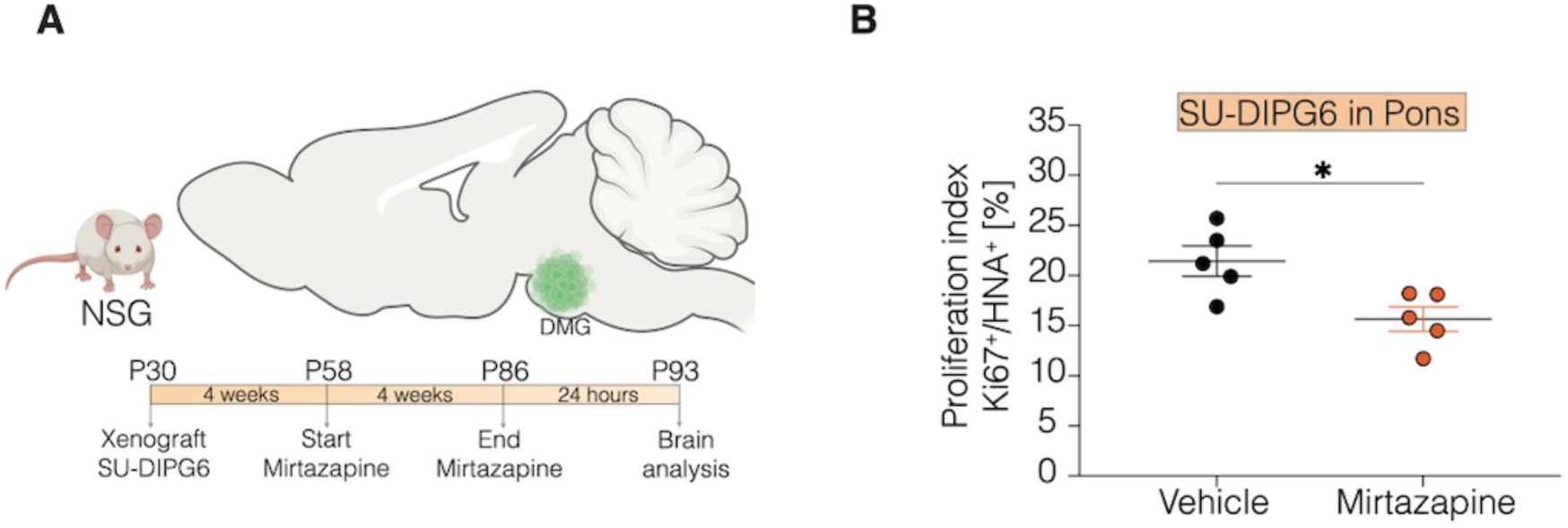
Mirtazapine reduces glioma cell proliferation. **A.** Schematic of the experimental paradigm for xenografting and chronic mirtazapine treatment. Four-week-old NSG mice (P28–30) were xenografted in the ventral pons with patient-derived DMG cells (SU-DIPG6). Daily intraperitoneal mirtazapine treatment (10 mg/kg body weight) was initiated four weeks post-xenograft and continued for four consecutive weeks. **B.** Proliferation index (Ki67⁺/HNA⁺) of ventral pontine xenografts in mice treated with the mirtazapine or vehicle control (*n* = 5 mice/group). Unpaired two-tailed Welch’s *t*-test; *p < 0.05. Data are presented as mean ± SEM.

## Materials and Methods

### Patient-derived Glioma Cell Cultures

IDH-wildtype glioblastoma and H3K27M diffuse midline glioma (DMG) cultures were established from patient-derived specimens with informed consent and in accordance with a protocol approved by the Stanford University Institutional Review Board (IRB). Patient-derived glioblastoma models included SF232, SF238, pcGBM6, GBM86r, GBM39 and UKE3 (a generous gift from the Brain Tumor Biology Laboratory Hamburg, Germany), while H3K27M DMG models included SU-DIPG6, SU-DIPG13fl, SU-DIPG13P, SU-DIPG17, SU-DIPG19, and SU-DIPG92. All cultures were routinely monitored for authenticity using short tandem repeat fingerprinting and tested for mycoplasma contamination. DMG cultures were maintained as neurospheres in serum-free medium consisting of DMEM (Invitrogen), Neurobasal(-A) (Invitrogen), B27(-A) (Invitrogen), heparin (2 ng ml−1), human bFGF (20 ng ml−1, Shenandoah Biotech), human bEGF (20 ng ml−1, Shenandoah Biotech), human PDGF-AA (10 ng ml−1, Shenandoah Biotech), and human PDGF-BB (10 ng ml−1, Shenandoah Biotech). For glioblastoma cultures, N2 supplement (Gibco) was added at a 1× concentration.

### MADR Glioma Model

The H3.3 K27M MADR tumor cell cultures were generated using the same technique as previously described(*21*, *43*). Briefly, Gt(ROSA)26Sortm4(ACTB-tdTomato,-EGFP)Luo/J and Gt(ROSA)26Sortm1.1(CAG-EGFP)Fsh/Mmjax mice (JAX Mice) were crossed with wild-type CD1 mice (Charles River) to produce heterozygous mice. Male and female embryos between E12.5 and E15.5 were subjected to in utero electroporation. Pregnant dams were individually housed, and pups remained with their mothers until P21 in the institutional animal facility (Stanford University). The MADR tumor cell line used here was generated by dissociating and sorting GFP+ tumor cells from female heterozygous mTmG mice. Subsequently, MADR cultures were maintained as neurospheres in serum-free medium composed of DMEM (Invitrogen), Neurobasal(-A) (Invitrogen), B27(-A) (Invitrogen), heparin (2 ng ml−1), human bFGF (20 ng ml−1) (Shenandoah Biotech), human bEGF (20 ng ml−1) (Shenandoah Biotech), human PDGF-AA (10 ng ml−1) (Shenandoah Biotech), insulin (Sigma-Aldrich), and 2-mercaptoethanol (Sigma-Aldrich).

### Animals Models

All animal experiments were conducted in accordance with protocols approved by the Stanford University Institutional Animal Care and Use Committee (IACUC) and performed in accordance with institutional guidelines. Animals were housed according to standard guidelines with unlimited access to water and food, under a 12 h light: 12 h dark cycle, a temperature of 21 °C and 60% humidity. For brain tumor allograft or xenograft experiments, the IACUC has a limit on indications of morbidity (as opposed to tumor volume). Under no circumstances did any of the experiments exceed the limits indicated and mice were immediately euthanized if they exhibited signs of neurological morbidity or if they lost 15% or more of their initial body weight.

Several mouse lines were used in this study. SERT-Cre mice (obtained from The Jackson Laboratory, strain #014554) were either used as a pure line with the genotype SERT-Cre_+/+_ or bred with homozygous transgenic Ai230 mice(*21*). Optogenetic experiments using allograft MADR models were performed on animals with the genotype SERT-Cre_+/wt_ x Ai230_flx/wt_, which enabled optogenetic control selectively for 5HT neurons in immunocompetent mice. To establish glioma xenografts from patient-derived glioma cells, immunodeficient NSG mice were used. To assess neuronal activity in the raphe nucleus, Fos2A-iCreER (TRAP2) mice (Jackson Laboratory, strain #030323) were crossed with B6.Cg-Gt(ROSA)26Sortm14(CAG-tdTomato)Hze/J (Ai14) mice (Jackson Laboratory, strain #007914). For fiber photometry experiments targeting GABAergic neurons of the raphe nucleus, B6J.129S6(FVB)-Slc32a1tm2(cre)Lowl/MwarJ (Vgat-ires-cre knock-in) mice (Jackson Laboratory, strain #028862) were used. All mice used were genotyped at postnatal day 10.

### Stereotactic Glioma Cell Implantation

Male and female mice were used in cohorts equally. For experiments conducted in immunocompetent mice, MADR cultures (H3.3 K27M MADR line 1; 200,000 cells per mouse) were allografted into either the M2 cortex, thalamus, or ventral pons. For experiments performed in immunodeficient NSG mice, patient-derived glioblastoma (SF232, UKE3, and GBM39-GCaMP6s; 300,000 cells per mouse) and DMG cultures (SU-DIPG6, SU-DIPG6-GCaMP6s, and SU-DIPG17; 300,000 cells per mouse, and SU-DIPG13P; 150,000 cells per mouse) were injected into the M2 cortex, thalamus, or ventral pons. A single-cell suspension of all cultures was prepared in sterile culture medium immediately prior to surgery. Animals at P28–P35 were anesthetized with 1–4% isoflurane and positioned in a stereotaxic apparatus. Under sterile conditions, the cranium was exposed via a midline incision, and a 31-gauge burr hole was made at the specified coordinates. For cortical M2 injections, coordinates were: AP = +1.0 mm (from bregma), ML = +0.5 mm, DV = –1.0 mm. For thalamus injections: AP = –1.0 mm (from bregma), ML = +0.8 mm, DV = –3.5 mm. For ventral pontine injections: AP = –0.8 mm (from lambda), ML = –1.0 mm, DV = –5.0 mm. Cells were injected using a 31-gauge Hamilton syringe at an infusion rate of 0.5 µl min−1 with a digital pump. At completion of infusion, the syringe needle was allowed to remain in place for a minimum of 5 minutes, then manually withdrawn. The wound was closed using 3M Vetbond (Thermo Fisher Scientific) and treated with Neo-Predef with Tetracaine Powder.

### Viral vectors and injections

Several AAVs were used in this study for *in vivo* experiments. The following AAVs were purchased from the Stanford Neuroscience Gene Vector and Virus Core: AAV-8-hSyn-GRAB_5-HT1.0 (GVVC-AAV-240), AAV-DJ EF1α-DIO-GCaMP 6m (GVVC-AAV-92), AAV-DJ-EF1α-DIO hChR2(H134R)-eYFP (GVVC-AAV-038), and AAV-DJ-EF1α-DIO eYFP (GVVC-AAV-013). Viral vectors were injected into the region of interest using a Hamilton Neurosyringe and Stoelting stereotaxic injector over a period of 5 minutes. All viruses were injected at a concentration of 2-8 × 10¹² infectious units per ml and were generally injected 0.3 mm below the ferrule placement (for DR, a DV of −3.4 mm; for MR, a DV of −4.5 mm). Additionally, to enable optogenetic stimulation of 5HT neurons in immunodeficient NSG mice engrafted with patient-derived glioma cultures, retro-orbital injections were performed using the previously validated viral constructs rAAVPHP.eB-Tph2::eYFP or rAAVPHP.eB-Tph2::ChRmine-eYFP(*44*), in collaboration with the Deisseroth Lab (Stanford, USA). For intravascular targeting of 5HT neurons, 4-week-old mice were injected retro-orbitally with 60 µL of virus solution (7 × 10_1_¹ vg/ml in 60 µL 0.9% NaCl per mouse). Mice were anesthetized with isoflurane and placed in a prone position on a heating pad to maintain body temperature. The injection site, located at the medial canthus of the eye, was cleaned with 70% ethanol. Using a 30-gauge needle, the virus solution was carefully injected into the retro-orbital sinus.

### Stereotactic Ferrule Placement

For all surgeries, mice of at least 4 weeks of age were anesthetized with 1–4% isoflurane and positioned in a stereotaxic apparatus. For optogenetic and fiber photometry experiments targeting 5HT neurons, optical fibers were implanted in either the dorsal raphe (DR) or median raphe (MR) nucleus. For the DR, fibers were implanted at the following coordinates relative to bregma: AP = −4.6 mm, ML = +0.0 mm, DV = −3.1 mm from skull. For MR targeting, coordinates were AP = −4.6 mm, ML = +0.0 mm, DV = −4.2 mm from skull. For terminal field stimulations of DR^5HT^ and MR^5HT^ projections in the ventral pontine and thalamic glioma microenvironments, a ferrule was placed 0.3 mm above the original tumor injection site (for thalamic allografts, at a DV of −3.2 mm; for ventral pontine allografts, at a DV of −4.7 mm). For optogenetic stimulation, ferrules with a 200 µm core diameter (MFC_200/240-0.22_4.5mm_MF2.5_FLT, Doric Lenses Inc.) were used; for fiber photometry, ferrules with a 400 µm core diameter (MFC_400/430-0.66_4mm_MF2.5_FLT, Doric Lenses Inc.) were used. All optical fibers were secured using stainless steel screws (thread size 00-90 × 1/16, Antrin Miniature Specialties) placed contralateral to the fiber, along with C&B Metabond and light-cured dental adhesive cement (Geristore A&B paste, DenMat).

### Optogenetic Stimulation of DR^5HT^ and MR^5HT^ Neurons

Optogenetic stimulations were performed one week after optic ferrule implantation. For experiments involving optogenetic stimulation of DR^5HT^ and MR^5HT^ neurons of SERT-Cre_+/wt_ x Ai230_flx/wt_ or NSG mice using the rAAVPHP.eB-Tph2::ChRmine-eYFP construct, freely moving animals were connected to a 595 nm high-power LED system via a monofiber patch cord to achieve stimulation of ChRmine. For terminal field stimulations of 5HT projections in the glioma microenvironment, a 473 nm diode-pumped solid-state laser system was employed to stimulate ChR2. Optogenetic stimulation, for both neuronal cell bodies and axon terminals, was performed at 20 Hz, ten 5 ms pulses of light delivery every 3 seconds. For optogenetic stimulation with ChR2, light was delivered at 10 mW from the tip of a 200-µm core optic fiber (NA = 0.22; Doric Lenses), corresponding to an irradiance of 17.47 mW/mm² across the DR and MR. For experiments with ChRmine, a power output of 1 mW at the fiber tip yielded an estimated irradiance of 1.2 mW/mm² within the nuclei. Activation of MR^5HT^ and DR^5HT^ neurons was verified by cFos immunostaining in an exploratory cohort perfused 90 minutes after stimulation. Potential light-induced heating damages were excluded by histological examination of the stimulated tissue after each session. Optogenetic stimulation session lasted for 30 minutes for 1-day stimulations and 10 minutes per day for 7-days stimulations. Animals were injected intraperitoneally with 40 mg/kg EdU (5-ethynyl-2’-deoxyuridine; Invitrogen, E10187) before the session, and 24 hours (for glioma cell proliferation analysis) after the start of the stimulation. In Ai230 models, light delivery results in stimulation. Thus, we designated the non-stimulated group as "mock," which received no light stimulation, and statistical analyses included mock stimulation of both the DR and MR.

### Immunohistochemistry

All mice were anesthetized with intraperitoneal injections of 2.5% Avertin (tribromoethanol; Sigma-Aldrich, T48402), and transcardially perfused with 20 ml 0.1M phosphate buffer saline (PBS). Brains were postfixed in 4% paraformaldehyde (PFA) overnight at 4°C before cryoprotection in 30% sucrose solution for 48 hours. For sectioning, brains were embedded in optimum cutting temperature (OCT; Tissue-Tek) and sectioned coronally at 40 µm or sagitally at 60 µm using a sliding microtome (Leica, HM450). For immunohistochemistry, brain sections were stained using the Click-iT EdU cell proliferation kit (Invitrogen, C10339 or C10337) according to manufacturer’s protocol. Tissue sections were then stained with antibodies following an incubation in blocking solution (3% normal donkey serum, 0.3% Triton X-100 in tris buffer saline) at room temperature for 30 minutes. Mouse anti-human nuclei clone 235-1 (1:100; Millipore, MAB1281), rabbit anti-TPH2 (1:500; Abcam, ab184505), chicken anti-green fluorescent protein (1:500; Aves Labs, GFP-1020), rabbit anti-RFP (1:500, Rockland, 600-401-379-RTU), rabbit anti-Ki67 (1:500; Abcam, ab15580) or rabbit anti-cfos (1:500; Synaptic Systems, 226 008) were diluted in 1% blocking solution (1% normal donkey serum in 0.3% Triton X-100 in TBS) and incubated overnight at 4°C. All antibodies have been validated in the literature for use in mouse immunohistochemistry. The following day, brain sections were rinsed three times in 1x TBS and incubated in secondary antibody solution for 2 hours at room temperature. All secondary antibodies were used at 1:500 concentration including Alexa 488 anti-rabbit, Alexa 488 anti-mouse, Alexa 488 anti-chicken, Alexa 594 anti-chicken, Alexa 647 anti-rabbit (all Jackson ImmunoResearch), and Alexa 555 anti-rabbit (Invitrogen). Sections were then rinsed three times in 1x TBS and mounted with ProLong Gold (Life Technologies, P36930).

### Confocal Microscopy and Quantification

All image analysis were performed by experimenters blinded to the experimental conditions or genotype. Cell quantification within allografted or xenografted tumors was conducted by acquiring z-stacks using a Zeiss LSM980 scanning confocal microscope (Carl Zeiss). A 1-in-6 series of coronal brain sections were selected, with 4 slices (40µm thickness) analyzed in the grafted brain area (RN, cortex, thalamus, or ventral pons). Brain tissue damaged during perfusion or tissue processing was excluded from histological analysis. Tumor cells were identified as GFP+ (MADR allografts) or HNA+ (patient-derived xenografts) and quantified in each field to determine the proliferation index, calculated as the percentage of GFP+ or HNA+ cells co-labeled with EdU. Quantification was performed using a 20x magnification field, with 6 fields selected from each of 4 brain slices per animal, in which all cells were quantified. The selection of the 6 fields was done systematically, covering both the tumor core and the tumor margin equally, with the needle injection site serving as the starting point. On average, this quantification method resulted in counting approximately 500 cells per animal.

### Survival Studies

For survival studies, mice were xenografted at 4 weeks of age with SU-DIPG13P cultures in the ventral pons or UKE3 in the M2 cortex. Following xenografting, mice were continuously monitored for signs of neurological deficits or health decline. Mice received daily intraperitoneal injections of pimavanserin (10 mg kg⁻¹; Tocris, #7667) formulated in 10% DMSO (Sigma), 10% Tween 80 (Sigma), and PBS, or vehicle control. The start and duration of treatment were adjusted according to the aggressiveness of each glioma model. Morbidity criteria included a 15% reduction in body weight or severe neurological motor deficits consistent with brain dysfunction (e.g., circling and barrel rolling for brainstem tumors; seizures and loss of gait for cortical tumors). Statistical analyses were performed using Kaplan–Meier survival curves with log-rank testing.

### Assessment of Raphe Nucleus Activity using TRAP2 x Ai14 mice

To label active neuronal populations in glioma-bearing and control-injected mice, Fos2A-iCreER (TRAP2) mice (Jackson Laboratory, #030323) were crossed with Ai14 reporter mice (B6.Cg- Gt(ROSA)26Sortm14(CAG-tdTomato)Hze/J; Jackson Laboratory, #007914) to generate TRAP2 x Ai14 offspring. At 4 weeks of age, mice were allografted with MADR cells or received control injections. Neuronal activity was captured 4 weeks post-implantation by administering 4-hydroxytamoxifen (4-OHT; 10 mg/kg, intraperitoneally) to induce Cre-mediated recombination and permanent tdTomato expression in neurons with recent Fos activation. Mice were perfused 6 hours after 4-OHT injection, and brains were processed as described above. Quantification was performed by counting the number of tdTomato-positive cells within the RN.

### Fiber Photometry

Fiber photometry was employed across different experimental paradigms. For all recording sessions, mice were individually housed in an empty box and shielded from external noise and stimuli. Prior to each recording session, mice were allowed to habituate for 10 minutes after being connected to the patch cord. Data acquisition was performed using Synapse software in conjunction with an RZ5P lock-in amplifier (Tucker-Davis Technologies). GCaMP6m, GCaMP6s, PsychLight2.0, and GRAB sensors were excited by frequency-modulated 465 and 405 nm LEDs (Doric Lenses). Optical signals were band-pass filtered through a fluorescence mini cube (Doric Lenses) and digitized at 6 kHz.

### Fiber Photometry Recordings from DR^5HT^ and MR^5HT^ Neurons

For recordings from DR^5HT^ and MR^5HT^ neurons, 4-week-old SERT-Cre_+/+_ mice were intracranially injected with AAV-DJ EF1α-DIO-GCaMP6m (virus titer: 8.0 × 10¹_2_ vg/ml). Three weeks post-injection, H3K27M-MADR cells or control solution (HBSS) were injected into either the thalamus or ventral pons. Three weeks after glioma cell implantation, an optical fiber (400 µm core diameter, NA 0.48; Doric Lenses) was implanted above the DR or MR. Following a one-week recovery period, recording sessions were initiated, and mice were recorded for 5 minutes per session.

### Fiber Photometry Recordings from Serotonin Release in the Glioma Microenvironment

For serotonin recordings in the glioma microenvironment, AAV-8-hSyn-GRAB_5-HT1.0 (virus titer: 7.1 × 10¹² vg/ml) was injected into 4-week-old SERT-Cre_+/+_ mice, positioned 0.5 mm above the ventral pontine tumor cell injection site. After allowing for viral expression, H3K27M-MADR cells or control solution were implanted into the ventral pons, with simultaneous placement of an optical fiber (400 µm core diameter, NA = 0.48; Doric Lenses) above the GRAB injection site. Fiber photometry recordings were performed one week after recovery, with mice recorded for 5 minutes per session.

### Fiber Photometry Recordings of Glioma Cells

We performed several experiments monitoring calcium signals from glioma cells transduced with either GCaMP6s or PsychLight2.0 (see below). These cell lines were used for recordings during simultaneous optogenetic stimulation of DR^5HT^ and MR^5HT^ neurons or following psilocybin injection. To monitor calcium signals from patient-derived glioblastoma or DMG cells during optogenetic stimulation, we utilized DMG (SU-DIPG6-GCaMP6s) and glioblastoma (GBM39-GCaMP6s) cell lines lentivirally transduced with GCaMP6s (pLV-ef1-GCaMP6s-P2A-nls-tdTomato). 4-week-old NSG mice were retro-orbitally injected with rAAVPHP.eB-Tph2::ChRmine-eYFP, as previously described, to enable optogenetic stimulation of 5HT neurons. Three weeks later, DMG cells were xenografted into the thalamus and glioblastoma cells were xenografted into cortex, with optical fiber placement (400 µm core diameter, NA = 0.48, Doric Lenses) above the implantation site and an additional fiber in the DR or MR for optogenetic stimulation. Recordings were initiated 6 weeks post-tumor xenograft. Mice were connected to both optogenetic and fiber photometry cables, with a 3-minute baseline recording followed by 3 minutes of optogenetic stimulation.

### Signal Processing and Data Analysis

Signal processing was performed using pMAT and custom MATLAB scripts (MathWorks). Raw fluorescence signals were de-bleached by fitting the data to either a mono-exponential or bi-exponential decay function. For normalization, fluorescence traces were Z-scored relative to a fixed baseline distribution, defined as the −10 to 0 s period immediately preceding the onset of the experimental intervention. The baseline distribution (mean and standard deviation) was pooled across all animals within each treatment or stimulation group for a given session. To visualize the Z-score data, a Locally Weighted Scatterplot Smoothing (LOESS) technique was applied to the time-series data. LOESS smoothing was performed with a smoothing fraction of 0.1 to generate a robust curve that effectively captures the underlying trends in the data while reducing noise. The area under the curve (AUC) was computed by integrating the Z-score signal over a defined time window following stimulus onset, using the trapezoidal rule to obtain a quantitative measure of the response magnitude.

### Transfection of Glioma Cells with PsychLight2.0

To enable recordings of PsychLight2.0 fluorescence in glioma cells (SF232 and SU-DIPG17), cells were transduced *in vitro* with PsychLight2.0(*45*), which was packaged into a lentiviral backbone. The PsychLight2.0 sequence was obtained from the published sequence of Addgene plasmid number 163910 (http://n2t.net/addgene:163910; RRID:Addgene_163910) and synthesized as a DNA fragment (Twist Biosciences). The plasmid pLenti-Ef1α-PsychLight2.0 was constructed by ligating the PsychLight2.0 fragment digested with BamHI and AscI into a pLenti-Ef1α backbone vector digested with the same restriction enzymes. The final construct was sequence-verified by whole plasmid sequencing. Additionally, lentivirus was produced for related experiments by the Stanford Neuroscience Gene Vector and Virus Core. Following packaging and titer optimization in HEK293T cells, glioma cells were transduced with LV-Ef1α-PsychLight2.0 for 72 hours using a transduction enhancer (TransDux MAX Lentivirus Transduction Enhancer, LV860A-1, System Biosciences). The lentiviral vector generated for this study is available upon request.

### Psilocybin Injections

Psilocybin powder (obtained from the National Institute of Drug Abuse) was diluted in 0.9% sterile saline to a concentration of 0.2 mg/mL. Psilocybin solution was then administered intraperitoneally at a volume of 10 mL/kg to achieve a dose of 2 mg/kg for all experiments. The head twitch response (rapid, side-to-side rotational head movements) was visually observed in all animals administered the drug.

### Collection of Conditioned Media from RN_5HT_ Neurons Following Ex Vivo Stimulation

SERT-Cre_+/wt_ x Ai230_flx/wt_ mice aged 4 weeks (P28–P30) were allografted with MADR cells or control solution (HBSS) into the ventral pons. Four weeks after tumor cell implantation, these mice were used to collect conditioned media from RN_5HT_ neurons. Brief exposure to isoflurane induced unconsciousness in the mice before immediate decapitation. Extracted brains (cerebrum) were inverted and placed in an oxygenated sucrose cutting solution, then sliced at 200µm to target the region of RN. Slices (n=3 per mouse) were transferred to ACSF and allowed to recover for 30 minutes at 37°C, followed by an additional 30 minutes at room temperature. The correct localization of the RN in each brain slice was verified by live microscopic identification of oScarlet-labeled 5HT neurons. After recovery, the slices were transferred to fresh ACSF and positioned under a red-light LED using a microscope objective. The optogenetic stimulation paradigm consisted of 20-Hz pulses of red light for 30 seconds on, followed by 90 seconds off, repeated over a period of 30 minutes. Conditioned medium from the surrounding area was collected and stored frozen at -80°C. Stimulated slices were postfixed in 4% paraformaldehyde (PFA) for 30 minutes before cryoprotection in 30% sucrose solution for 48 hours. Successful stimulation of RN_5HT_ neurons in each area was validated through cFos staining.

### Nano Liquid Chromatography–Tandem Mass Spectrometry

NanoLC-MS/MS and data analysis was performed by Applied Biomics (www.appliedbiomics.com/) under the LCMSMS protocol number MOMI241002.

#### Sample preparation

Media samples were concentrated to 4 mg/mL using spin columns with a 3 KDa molecular weight cut-off (MWCO), then buffer-exchanged into 50 mM ammonium bicarbonate. Protein concentration was determined using the Bio-Rad Protein Assay Kit II (#500-0002) according to the manufacturer’s instructions. For each sample, 30 µg of protein was used. DTT was added to a final concentration of 10 mM, and samples were incubated at 60 °C for 30 minutes, followed by cooling to room temperature. Iodoacetamide was then added to a final concentration of 10 mM, and samples were incubated in the dark at room temperature for 30 minutes. Proteins were subsequently digested overnight at 37 °C using Trypsin/Lys-C Mix, Mass Spec Grade (Promega). *nanoLC-MS/MS* NanoLC separation was performed using a Thermo Fisher Ultimate 3000 system (Milford, MA). Mobile phases consisted of solvent A: 0.1% formic acid (v/v) in water, and solvent B: 0.1% formic acid (v/v) in 90% acetonitrile. Tryptic peptides were first loaded onto a µPrecolumn Cartridge (serving as a trap column for on-line desalting), followed by separation on a reverse-phase analytical HPLC column (Thermo Fisher Scientific, ES900). The chromatographic gradient was as follows: 2% to 10% solvent B over 10 minutes, 10% to 30% B over 85 minutes, followed by a wash at 99% B for 10 minutes and re-equilibration at 4% B for 5 minutes. Mass spectrometry was conducted on an Orbitrap Exploris (Thermo Fisher Scientific). MS1 scans were acquired at 80,000 resolution over a scan range of m/z 300–2000, with 100% AGC target and maximum injection time set to auto. MS/MS scans were performed at 30,000 resolution with a 200% AGC target, auto maximum injection time, 30% normalized HCD collision energy, and an intensity threshold of 1.0e4.

### Database search

Data analysis was conducted using Proteome Discoverer 2.5 (Thermo Fisher Scientific). MS and MS/MS spectra were searched against the SwissProt Mus musculus database, containing all protein entries as of October 2024. Searches were performed using the Sequest HT algorithm with the following parameters: enzyme specificity set to Trypsin, minimum peptide length of 5, maximum peptide length of 144, precursor mass tolerance of 10 ppm, and fragment mass tolerance of 0.02 Da. Dynamic modifications included oxidation of methionine (Oxidation-M), N-terminal acetylation, and N-terminal methionine loss. Carbamidomethylation of cysteine (Carbamidomethyl-C) was set as a static modification. Label-free quantification of individual and grouped protein abundances was performed using the Minora Feature Detector. For both the processing and consensus workflows, results were filtered to a maximum false discovery rate (FDR) of 1%. Protein abundance values and ratios were normalized based on total peptide amount.

### CRISPR Deletion of HTR2A

Patient-derived DMG cells (SU-DIPG17) were cultured in complete growth media at 37°C in a 5% CO2 incubator as described above. Cells were passaged at 80% confluency and maintained in optimal conditions. To generate a Cas9-expressing cell line, we used a lentiviral vector (pB-TRE3G-spCas9-Ubc-rtTA3-IRES-BlastR, Addgene #195506). For the generation of a Cas9-expressing DMG cell line (SU-DIPG17-Cas9), we used the inducible PiggyBac vector Pb-TRE3G-SpCas9-IRES-Blast (Addgene #195506), The transposition system for the inducible Cas9 vector was performed using the PiggyBac transposase system. DMG cells were transfected with 2.5 µg of the Pb-TRE3G-SpCas9-IRES-Blast vector and 0.5 µg of the PiggyBac transposase plasmid using FuGENE HD according to the manufacturer’s protocol. After 6 hours, the cells were provided with fresh media and allowed to recover for 6 days. Transfected cells were selected with 5 µg/mL blasticidin (Thermo Fisher Scientific). Successfully transduced cells, expressing both Cas9 and the blasticidin resistance marker, were expanded. Cas9 expression was confirmed by Western blotting. For CRISPR-mediated knockout of HTR2A, guide RNA (gRNA) plasmids were designed to target the top three coding exons of HTR2A (gRNA sequence: CGATTCGAAGGTCTTTAAGG; chr13[46835602], GGTCGATCCATAGGGAGCCA; chr13[46835363], TAGCGACGGAGTGAATGAAA; chr13[46834873]; Thermo Fisher Scientific) using the TrueDesign Genome Editor (Invitrogen). DMG cells were passaged at 80% confluency and filtered into a single-cell suspension before electroporation to ensure optimal cell density. The cells (n=100.000) were electroporated with a total volume of 100uL OPTI-MEM (Gibco) containing gRNAs targeting HTR2A. Electroporation was performed using the NEPA21 Electroporator (NEPA GENE) and the CU500 Cuvette Chamber (Bulldog Bio) under the following conditions: 300V pulse voltage, 50 ms pulse interval, 1 pulse, and a 5 ms pulse time. After electroporation, cells were immediately transferred into 2 mL of complete growth media and incubated at 37°C for 24 hours to recover. Twenty-four hours post-electroporation, the cells were selected with blasticidin (Thermo Fisher Scientific).

### EdU Proliferation Assay In Vitro

EdU staining of glioma monocultures was performed on glass coverslips in 96-well plates which were pre-coated with poly-l-lysine (Sigma) and laminin (Thermo Fisher Scientific). Neurosphere cultures were dissociated with TrypLE and plated onto coated coverslips with growth factor-depleted medium. Serotonin (1µM to 10µM, Tocris), LSD (100mM), DOI (100mM), and vehicle (DMSO) were added for specified times with 4 µM EdU. After 24 h the cells were fixed with 4% PFA in PBS for 20 min and then stained using the Click-iT EdU kit and protocol (Invitrogen) and mounted using Prolong Gold mounting medium (Life Technologies). Confocal images were acquired on a Zeiss LSM980 using Zen 2011 v8.1. Proliferation index was determined by quantifying the fraction of EdU-labelled cells divided by DAPI-labelled cells using confocal microscopy at 20× magnification. Per coverslip, 6 fields were imaged and quantified, including all DAPI_+_ cells within each field. The fields were selected systematically, with 3 fields from the center and 3 fields from the margins of the coverslip. Using this method, an average of 1000 cells per coverslip were counted. Quantification of images was performed by a blinded investigator.

### Measurement of Serotonin In Vitro

The concentration of serotonin (Eagle Biosciences, #EA630/96) in media from various patient-derived glioma cultures was measured using ELISA according to the manufacturer’s instructions. Optical density was measured at 450 nm using an absorbance microplate reader (SpectraMax M3, Molecular Devices). Serotonin concentrations were calculated as µg/mL based on the standard curve. To confirm the specificity of the assay, medium and lysis buffer without protein extract were included as negative controls.

### Forced Swim Test

The forced swim test was employed to assess depressive-like behavior potentially indicative of altered activity within the raphe nucleus in ventral pontine glioma-bearing mice compared to control-injected animals. SERT-Cre_+/+_ mice underwent a pre-test at 4 weeks of age, followed by ventral pontine glioma allografting or control injection (HBSS) 24 hours later. Forced swim testing began 7 days post-surgery and was repeated weekly on the same days throughout the study. Mice were habituated to the testing room for at least 15 minutes prior to each session. Each animal was placed in a transparent plastic cylinder (25 cm height × 18.5 cm diameter) filled with water (23–25 °C) to a depth of 14 cm, ensuring that the hind limbs could not touch the bottom. Mice were tested for 6 minutes while behavior was recorded using a camera positioned in front of the cylinder. The initial 2 minutes were considered an acclimation period; immobility was quantified during the final 4 minutes. Immobility was defined as the absence of active movement, except for minimal motions required to keep the head above water. All mice completed the full experimental paradigm through the final session at 4 weeks post–tumor cell implantation, ruling out reduced motor function in tumor-bearing animals. Immobility analysis was performed by two independent experimenters blinded to genotype and condition, and immobility time was calculated as the average of both assessments. A two-day testing paradigm was applied. Water was replaced between animals to prevent olfactory cue bias. After each session, mice were dried before being returned to their home cages.

### Analysis of single-cell RNA sequencing datasets

We analyzed single-cell RNA-sequencing (scRNA-seq) data from 18 primary patient samples with H3K27M DMG, comprising a total of 5,111 malignant cells, generated using SMART-seq reported by Liu et al., to investigate the expression of genes in the 5-HTR family. Data processing for diffuse midline glioma (DMG), including the definition of cell states and cellular hierarchies, followed the procedures described in the original publication(*41*). In addition to evaluating gene expression across defined cell states, we assessed the expression of 5HTR family genes across tumor locations, including pontine, spinal, and thalamic regions. Finally, co-expression patterns of HTR2A and NTRK2 were visualized using the R package Nebulosa (v1.18.0), specifically employing the plot_density function to generate UMAP representations.

### Spatial transcriptomics analysis of human glioma samples

Spatial transcriptomics data were generated as described in Ravi et al. (*42*). Briefly, Spatial transcriptomics experiments were conducted on adult IDH wild-type glioblastoma and pediatric H3K27M diffuse midline glioma tumors using the 10X Visium Spatial Gene Expression kit, following the manufacturer’s protocol for tissue optimization and library preparation. Fresh tissue samples were collected immediately after biopsy, embedded in Tissue-Tek O.C.T. Compound, and snap-frozen in isopentane cooled with liquid nitrogen. The tissue was stored at −80 °C until further processing. Ten-micrometer tissue sections per sample were lysed using TriZol, and RNA was isolated using the PicoPure RNA Isolation Kit. RNA integrity was assessed using a Fragment Analyzer, and only samples with an RNA integrity number (RIN) greater than 7 were used for further analysis. For the spatial gene expression protocol, 10 µm tissue sections were mounted on spatially barcoded glass slides with poly-T reverse transcription primers, one per array, and fixed with 100% methanol. H&E staining and brightfield imaging were performed at 10X magnification, followed by permeabilization to capture mRNA on the primers. cDNA was generated using template switch oligos and amplified with KAPA SYBR FAST qPCR Master Mix. After size selection using SPRIselect reagent, quality control was performed with a Fragment Analyzer. The cDNA was further optimized for sequencing using Illumina NextSeq, with unique indexes and Illumina primers added. Final libraries were quantified and sequenced on the Illumina NextSeq 550 platform, using paired-end sequencing with 28 cycles for reading 1, 10 cycles per index, and 120 cycles for read 2 on a NextSeq 500/550 High Output Kit v2.5 (Illumina, 20024907).

To evaluate cell type locations in the Visium spatial transcriptomics data, we used the Cell2location model with the abovementioned scRNA-seq data as a reference. We setup the Cell2location model by configuring an AnnData object, defining parameters such as the number of cells per location and detection sensitivity. The model was trained on a graphics processing unit for 500 iterations. After training, we extracted the posterior distribution of cell type proportions, sampling 1,000 times for precise estimation across the spatial framework. Median estimates of cell type abundance were saved in the AnnData object. Finally, we incorporated the cell type abundance data back into the SPATA object using the addFeature function in SPATA2, allowing for further analysis steps within the spatial data framework(*46*).

Spatial enrichment scores for HTR2A and NTRK2 was calculated for individual Visium spots as described above. Deconvoluted cell type abundance from the Cell2location algorithm were used for cell type specific correlation analysis with spatial enrichment scores for both genes.

Pearson’s r to determine correlation. Gene set enrichment analysis was performed using the SPATA2 package with all gene sets included in standard workflow.

## Statistical analysis

To determine the appropriate sample size for *in vivo* proliferation studies, a power analysis was conducted using an unpaired t-test to compare the means of two independent groups. The significance level was set to 0.05 (two-tailed), with a target power of 0.8. Based on pilot data for our allograft and xenograft models in combination with optogenetic stimulation, the power analysis indicated that a sample size of 4 animals per group would be required to achieve 80% power. The achieved power for this sample size was calculated to be 0.939. Gaussian distribution was confirmed using the Shapiro-Wilk test. Parametric data were analyzed with unpaired two-tailed Welch’s t-tests or one-way ANOVA with Tukey’s post hoc tests. Significance was set at p < 0.05. The used statistical test is indicated in the figure legend. GraphPad Prism 10 was used for statistical analyses and data illustrations.

